# Foundational genomic resources for date palm: A gap-free, telomere-to-telomere phased assembly of Ajwa and 19 high-quality genome assemblies of *Phoenix dactylifera*

**DOI:** 10.64898/2025.12.30.696066

**Authors:** Mirko Celii, Noor Al-Bader, Nahed Mohammed, Yong Zhou, Alice Fornasiero, Maria Navarrete Rodriguez, Raghad Shuwaikan, Umair Toor, Jun Hong, Victor Llaca, Kevin Fengler, Dal-Hoe Koo, Muriel Gros-Balthazard, Vincent Battesti, Thiago Capote, Michael Purugganan, Joel A. Malek, Ikram Blilou, Jesse Poland, Rod A. Wing

## Abstract

*Phoenix dactylifera* L. is an economically, nutritionally, and culturally important fruit crop in the arid and semi-arid regions of the Middle East and North Africa. Here, we present a gap-free, telomere-to-telomere reference genome of the variety Ajwa, along with 19 additional high-quality assemblies (18 female and 1 male). These assemblies reveal novel chromosomal structures validated through cytogenetics, Hi-C, optical mapping, and synteny analyses with other palm genomes. Chromosome names were revised based on average lengths across all sequenced genomes. The Sex Determination Region (SDR) on chromosome 14 was confirmed through male-specific k-mer analysis, spanning approximately 14.7 Mb. Nucleolar organizing regions (NORs) were localized on chromosome 10, where a large 45S rDNA locus displayed unique repeat spacer motifs containing transposon-like sequences. In some accessions, a second NOR was identified on the female sex chromosome. This collection of date palm assemblies, anchored by the Ajwa reference genome, provides a critical resource for advancing breeding strategies aimed at enhancing the genetic resilience and productivity of date palm.

## Background & Summary

Date palm is one of the most important fruit-bearing crops cultivated across hot and arid environments, primarily spanning from Morocco to South Asia, and is especially associated with oasis agroecosystems ^1^. It has remarkable adaptability and resistance to environmental stresses such as heat and drought, which makes it a valuable model for genetic studies of plants in arid regions. Despite its strategic importance, high-quality genomic resources for date palms remain limited. To date, two genome assemblies have been published: Khalas PDK50 ^2,3^ and Barhee BC4^4^. However, both assemblies present significant limitations, due primarily to the sequencing technology employed. For example, the Khalas genome, while being highly contiguous, contains an exceptionally high number of gaps (10,972). In contrast, the Barhee genome, although featuring relatively fewer gaps (317), could not be fully anchored to the Khalas chromosomes, leaving ∼50% of its genome unplaced (i.e. 385 Mb out of 772 Mb). To address this lack of genomic resources we present here the generation of a gap-free, telomere-to-telomere (T2T) phased assembly of the variety Ajwa. The Ajwa variety (ʿajwah) was selected due to its cultural and commercial importance for the Kingdom of Saudi Arabia (KSA). The voucher plant was provided by the al-Dabta farm in Medina, KSA (https://aldabtafarm.com/). This new assembly features an updated chromosome structure and refined genome size, which we propose to serve as the reference standard genome for date palm genomics. In addition, we present 18 additional high-quality female genome assemblies from important varieties that are grown in the KSA, as well as a high-quality male genome assembly featuring a gap-free and T2T-assembled sex chromosome.

## Methods and Results

### Ajwa Genome Karyotyping with Fluorescence *in situ* hybridization (FISH)

To confirm previous karyotyping studies of *Phoenix dactylifera* L.^2,5,6,7^ and map the location of 45S rDNA, we performed fluorescence *in situ* hybridization (FISH) following the methods described by Koo *et al*.^8^ with minor modifications. For FISH, we obtained root tips from actively growing young roots, which were obtained by growing seedlings from seeds from Ajwa fruits. To germinate the seeds, date seeds were wrapped in a wet paper towel and kept in a dark place for approximately 21 days until sprouting, then transplanted into pots. Mitotic chromosome spreads were prepared from root tips of actively growing plants. Root tips were treated with nitrous oxide for 1.5 h, fixed overnight in ethanol: glacial acetic acid (3:1), and then squashed in 45% acetic acid. The preparations were stored at –70°C until use. Arabidopsis telomeric DNA (TTTAGGGn) and maize 45S rDNA clones^9^ were used as probes to identify the telomeric regions and nucleolar organizer region (NOR) of date palm chromosomes, respectively. The hybridization mixture (20 µl) contained 300 ng of each probe, 50% formamide, 2×SSC, and 10% dextran sulfate. Slides were denatured and hybridized for 18 h at 37°C. After post-hybridization washes, digoxigenin- and biotin-labeled probes were detected with rhodamine-conjugated anti-digoxigenin (Roche) and Alexa Fluor 488–conjugated streptavidin (Invitrogen), respectively. Chromosomes were counterstained with 4′,6-diamidino-2-phenylindole (DAPI) in Vectashield antifade solution (Vector Laboratories, Burlingame, CA). Images were captured using a Zeiss Axioplan 2 microscope (Carl Zeiss Microscopy LLC, Thornwood, NY) equipped with a cooled CCD camera (CoolSNAP HQ2, Photometrics, Tucson, AZ) and AxioVision 4.8 software. Final image contrast was adjusted using Adobe Photoshop 2024 (Adobe Systems Inc., San Jose, CA).

These FISH experiments displayed a diploid set of 36 chromosomes with two 45S rDNA-rich chromosomes, one for each haplotype (Figure 1) confirming both a chromosome number of 18 and the presence of a single-chromosome Nucleolar Organizing Region (NOR).

**Fig. 1.**
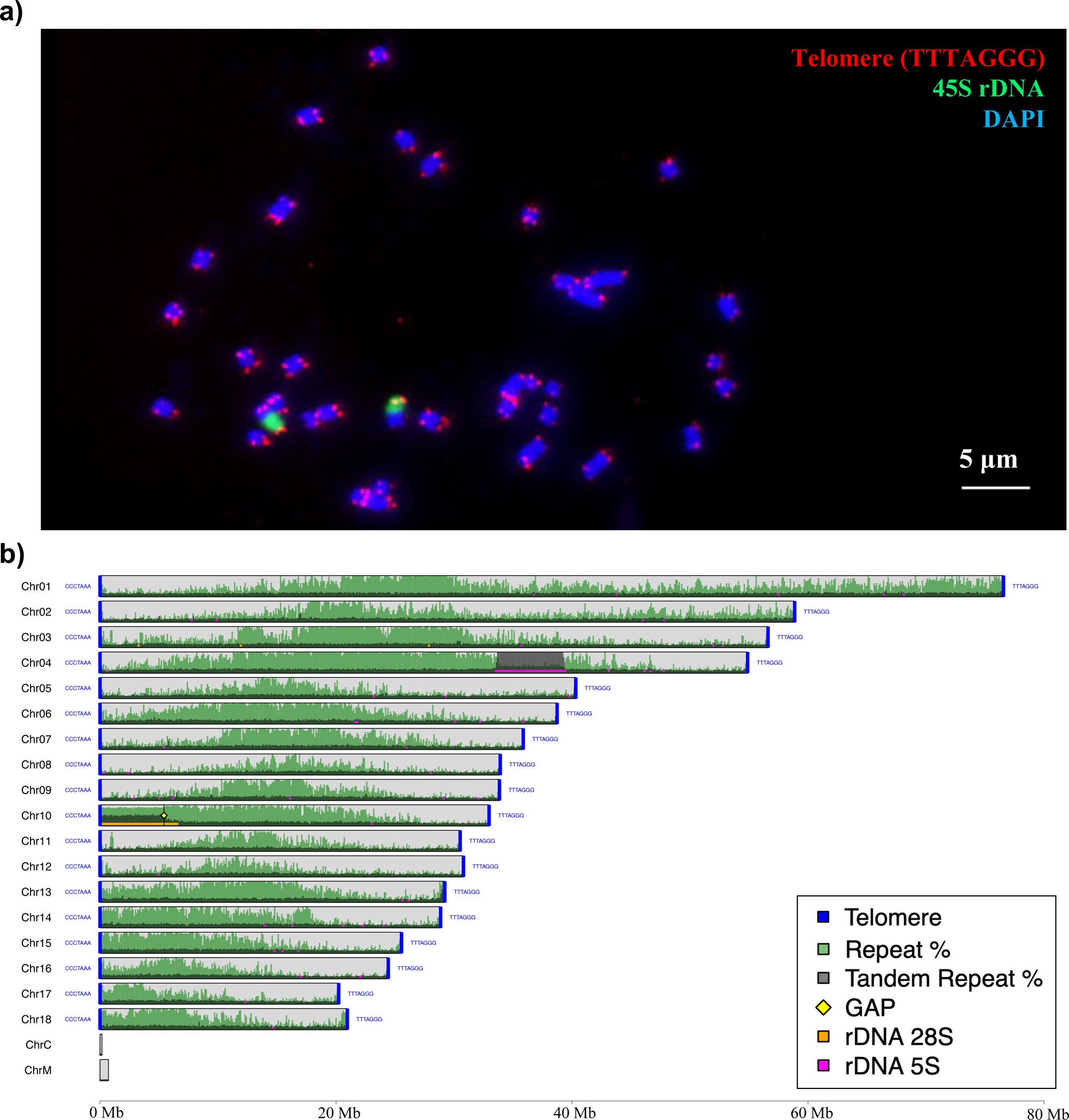
a) Fiber FISH of Ajwa seedlings using probes for Arabidopsis telomeric DNA (TTTAGGGn) and maize 45S rDNA clones^9^. b) Total repeat and tandem repeat profile of the Ajwa genome (Haplotype 1) shown as percentage in 100 Kb windows with the locations of telomeres, 5S rDNA and 45S rDNA, generated with GS-viewer repeat-profile.

### High Molecular Weight (HMW) DNA Extraction and Quality Control

Approximately 2 grams of flash frozen leaf tissue from a voucher plant for each date palm variety was ground into a fine powder using a mortar and pestle under liquid nitrogen (LN). HMW DNA was extracted using the Qiagen Genomic-tip 500/G Kit® (Qiagen, Germany), following the manufacturer’s protocol. DNA concentration and purity were initially assessed using a Qubit Fluorometric Quantification system (Invitrogen, USA) and a NanoDrop spectrophotometer (Thermo Fisher Scientific, USA). To evaluate DNA integrity and restriction accessibility, samples were digested with *Eco*RI and *Hind*III restriction enzymes, followed by gel electrophoresis. DNA fragment size distributions were then estimated using a Femto Pulse System (Agilent Technologies, USA) prior to library preparation.

### HMW DNA, PacBio Library Preparation and Sequencing

All PacBio libraries were generated and sequenced at KAUST Bioscience Core Laboratory (BCL), except for the first three Ajwa libraries (Supplementary Table 1a), replicates 1, 2 and 3) which were generated and sequenced at Arizona Genomic Institute (Tucson, Arizona, USA) as follows: DNA was sheared to an appropriate size range (20.5 kb) using a g-Tube (Covaris) with 4700 rpm centrifuge speed (Eppendorf® MiniSpin plus), followed by AMPure PB bead purification. The library was constructed following manufacturers’ protocols using a SMRTbell Express Template Prep kit 2.0. The final library was size selected on a BluePippin electrophoresis system (Sage Science) using the S1 marker with an 11-25 kb size-selection range. The recovered final library was quantified with a Qubit HS kit (Invitrogen) and size checked with a Femto Pulse System (Agilent). The final library was prepared for sequencing with a PacBio Binding kit 2.2, a Sequel II Sequencing kit 2.0, and loaded into three 8M SMRT cells, and sequenced in the CCS mode for 30 hours. In total, 8.13 million reads were generated with an average read length of X, yielding 156 GB of raw data, equivalent to approximately 110× per haplotype.

At the KAUST Bioscience Core Lab, HMW DNA was sheared to an average fragment size of ∼10–30 kb using a Megaruptor 3 (Diagenode, USA). Library preparation was performed using a SMRTbell Prep Kit 3.0 (PacBio), following manufacturer’s instructions. Size selection of the final library was carried out with a PippinHT electrophoresis system (Sage Science, USA) with a selection of fragments in the 10–25 kb range. Library concentration was measured using a Qubit Fluorometer High-Sensitivity Kit (Invitrogen), and fragment size distribution was assessed with a Femto Pulse System (Agilent Technologies, USA). Sequencing primer binding was performed using the Sequel II Binding Kit 3.2, and libraries were sequenced on 8M SMRT Cells in Circular Consensus Sequencing (CCS) mode using the PacBio Sequel II Sequencing Kit 2.0, with 30-hour runtimes on the Sequel II platform.

### Omni-C and Hi-C Library Preparation and Sequencing

For Ajwa, Omni-C libraries were generated by Dovetail Genomics (Scotts Valley, CA, USA). Chromatin was fixed in place with formaldehyde in nuclei and then extracted. Fixed chromatin was digested with DNAse I, chromatin ends were repaired and ligated to a biotinylated bridge adapter followed by proximity ligation of adapter containing ends. After proximity ligation, crosslinks were reversed and the DNA purified. Purified DNA was treated to remove biotin that was not internal to ligated fragments. Sequencing libraries were generated using a NEBNext® Ultra library preparation kit (New England Biolabs, Ipswich, Massachusetts, United States) and Illumina-compatible adapters. Biotin-containing fragments were isolated using streptavidin beads before PCR enrichment of each library. The library was sequenced on an Illumina HiSeq X platform to produce approximately 30x sequence coverage. For all other 19 date palm samples, Hi-C libraries were generated by Corteva Agriscience (Indianapolis, Indiana, USA), using the Proximo system (Phase Genomics), following the manufacturer’s recommended protocol. Proximo Hi-C libraries were pooled and sequenced in an Illumina NovaSeq system using PE150 as per manufacturer standards to a genome coverage of 43x. For each library, adapter sequences and low-quality bases were removed using Trimmomatic^10^, and quality control was assessed with FastQC^11^ (Supplementary Table 1b).

### RNA Extraction and Quality Assessment

Total RNA was extracted from leaf, root, and flower tissue of the Ajwa voucher plant, with each tissue type ground separately in liquid nitrogen using a mortar and pestle. RNA was isolated using a Maxwell® RSC Plant RNA Kit on the Maxwell® RSC 48 Instrument (Promega), following the manufacturer’s protocol. RNA concentration was determined using a Qubit Fluorometer (Invitrogen), and RNA integrity was evaluated using either the Agilent 4200 TapeStation System or the Agilent 2100 Bioanalyzer System, with RNA Integrity Number (RIN) values used to assess sample quality.

### Short- and Long-Read RNA Library Preparation and Sequencing

For the Ajwa date palm, RNA sequencing libraries were prepared using both short- and long-read sequencing technologies following manufacturer-recommended protocols. For short-read sequencing, 1 µg of total RNA was used to construct stranded total RNA libraries with the TruSeq Stranded Total RNA with Ribo-Zero Plant Kit (Illumina), using SuperScript IV Reverse Transcriptase (Invitrogen) for first-strand cDNA synthesis. Paired-end sequencing (2 × 150 bp) was performed on the Illumina NovaSeq 6000 platform. For long-read sequencing, full-length cDNA libraries (Iso-Seq) were prepared using an Iso-Seq Express Oligo Kit (PacBio, 101-737-500), following PacBio’s protocol (PN 102-396-000 REV02 APR2022). First-strand cDNA synthesis and amplification were carried out using 300 ng of total RNA and the NEBNext® Single Cell/Low Input cDNA Synthesis & Amplification Module (NEB E6421S). Barcoded forward and reverse cDNA primers (IDT) were used during amplification. Final Iso-Seq libraries were constructed using 500 ng of cDNA with the PacBio SMRTbell Prep Kit 3.0, which included cDNA repair, A-tailing, adapter ligation, nuclease treatment, and SMRTbell cleanup steps. Libraries were quantified using a Qubit Fluorometer (Invitrogen) and sized on an Agilent Bioanalyzer using the High Sensitivity DNA Kit. Sequencing-ready libraries were prepared according to SMRT Link Sample Setup and Run Design protocols. Sequencing complexes were bound using the Sequel II Binding Kit 3.1 and loaded onto 8M SMRT Cells. Sequencing was performed on a PacBio Sequel II System in CCS mode using the Sequel II Sequencing Kit 2.0, with 24-hour runtimes, obtaining a minimum of 100 million PE reads for a coverage of at least 44x (Supplementary Table 1b).

### Bionano Optical Genome Maps

HMW DNA for all individuals was isolated as described in Rabanal *et al.*^12^, with minor modifications as follows. Approximately 0.5 g of fast frozen healthy, young leaves were ground in liquid nitrogen and placed in ice-cold Bionano Homogenization buffer without a fixing step. Nuclei and cell debris were pelleted by centrifugation at 2,500 g before and after low-speed centrifugation steps to remove cell debris. Optical mapping was performed using the direct labeling and stain approach (Bionano Genomics; DLS), as described by Ou *et al.*^13^. Data processing, optical map assembly, hybrid scaffold construction and visualization were performed using the Bionano Solve (Version 3.4) and Bionano Access (v12.5.0) software packages (https://bionanogenomics.com).

### Heterozygosity Estimation

To estimate heterozygosity, PacBio HiFi reads were analyzed using Jellyfish v2.3.0^14^ and GenomeScope v2.0^15^, to generate counts of observed k-mers. This analysis, using a k-mer length of 51, revealed a bimodal distribution (Supplementary Figure 1), which is a typical feature of diploid heterozygous genomes.

The k-mer analysis estimated a high heterozygosity level of ∼0.7%, consistent with previous reports ^16,17^.

### Ajwa Phased Genome Assembly

To generate highly contiguous and haplotype resolved reference genome assemblies for the Ajwa variety, Pacbio high-fidelity reads were assembled using Hifiasm v0.19.0^18^ including Omni-C data (98.5 million paired-end reads) from Dovetail Genomics (Supplementary Tables 1a-b). Contaminant sequences were removed from the assembly using Kraken2 ^19^, along with contigs smaller than 70 kb. Contigs belonging to chloroplast and mitochondrial genomes were identified using Oatk^20^.

The resulting nuclear assembly size for haplotype 1 (Hap1) was 730 Mb, and 693 Mb for haplotype 2 (Hap2). Both values were notably smaller than those reported for the first two published date palm genomes, Khalas (854 Mb) and Barhee (772 Mb), as well as the 843 Mb genome size estimated by flow cytometry^21^. N50 and L50 values were 30.9 Mb and 9 for Hap1, and 34.1 Mb and 8 for Hap2, indicative of high-quality, haplotype-specific assemblies (Supplementary Table 2a-b).

### Ajwa Phased Scaffolding

Contigs derived from the two haplotypes were merged into a single assembly and subsequently scaffolded onto the optical maps using Bionano Solve v3.7 (https://bionanogenomics.com/) with default parameters, generating a hybrid scaffold. Resulting gaps were manually filled with Sequencher^22^ using corrected reads generated by Hifiasm. Tandem Repeat Finder^23^ was run to search for telomeric and centromeric repeats. Canonical telomeric sequences (TTTAGGGn) were identified at both termini of all chromosomes, except one, whereas no centromeric/pericentromeric tandem repeat patterns were detected. The final phased genome consisted of two sets of 18 chromosomes, with 4 gaps only (three within the NORs) and a single missing telomere (Supplementary Figure 2).

All 18 Ajwa chromosomes for each haplotype exhibited a perfect one-to-one correspondence with the 18 Bionano hybrid scaffolds with the sole exception of the NOR region on Chr10. As expected in this complex repetitive region, multiple unplaced contigs consisting of 45S rDNA repeats showed similar and overlapping patterns with the NOR. To further validate this structure, we applied the Hi-C scaffolding tool YAHS^24^ which similarly produced 18 Hi-C scaffolds with a direct correspondence to the Ajwa chromosomes (Supplementary Figure 3a-b). For each chromosome pair, we designated haplotype 1 (Hap1) the contig/pseudo-molecule with the optimal features (i.e. presence of both telomeres, absence or lowest number of gaps, higher length). Although no centromeric tandem repeats were detected, putative centromere locations were inferred based on regions with the highest repeat density, and chromosomes were oriented with the short arm positioned at the chromosome start (position zero). Similarly, organellar contigs were scaffolded on Bionano optical maps, resulting in gap-free genomes of the chloroplast (158.5 kb) and mitochondrion (715.1 kb) (Figure 1b, Table 1).

**Table 1.** Chromosome-level assembly statistics for the Ajwa genome and their correspondence to linkage groups in the Khalas PDK50 genetic map.

| Ajwa chromosome | Khalas Linkage Group | Ajwa Chromosome length | # gaps | notes |
| --- | --- | --- | --- | --- |
| **Chr01** | **LG04**<br>**LG16** | **76557843** | 0 |  |
| Chr02 | LG01 | 58863253 | 0 |  |
| Chr03 | LG02 | 56608022 | 0 |  |
| Chr04 | LG14 | 54878859 | 0 | rDNA 5S |
| Chr05 | LG08 | 40314578 | 0 |  |
| Chr06 | LG05 | 38739638 | 0 |  |
| Chr07 | LG06 | 35867188 | 0 |  |
| Chr08 | LG03 | 33913642 | 0 |  |
| **Chr09** | **LG11** | **33847661** | 0 |  |
| Chr10 | LG18 | 32967886 | 1 | rDNA 45S |
| Chr11 | LG07 | 30518302 | 0 |  |
| Chr12 | LG09 | 30798259 | 0 |  |
| Chr13 | LG10 | 29217882 | 0 |  |
| **Chr14** | **LG12** | **28862253** | **0** | Sex Chr |
| Chr15 | LG15 | 25550550 | 0 |  |
| Chr16 | LG17 | 24427294 | 0 |  |
| Chr17 | LG13 | 20226707 | 0 |  |
| **Chr18** | **LG11** | **20956597** | 0 |  |
| ChrC | na | 158463 | 0 |  |
| ChrM | na | 715125 | 0 |  |

### NOR Assembly and Size Estimation

The vast majority of remaining unscaffolded contigs in both haplotype assemblies consisted entirely of 45S rDNA copies, totaling approximately ∼38 Mb per haplotype. These contigs exhibited an average weighted coverage of ∼46x, approximately 2.35-fold lower than the rest of the haplotigs (∼107x), suggesting an overestimation of their total rDNA length. By dividing rDNA total size of ∼38 Mb by the coverage ratio of 2.35, we can estimate a more reliable unscaffolded 45S rDNA content of ∼16 Mb. However, due to the highly repetitive nature of these sequences and the lack of unique flanking regions, they could not be manually incorporated into the NOR regions and were thus excluded from the final genome assembly.

Of note, we observed a contig with 45S rDNA repeats directly adjacent to the telomeric sequences. This genomic structure of the rDNA ending at the telomere was confirmed by cytogenetics as shown in Figure 1a. This telomeric 45S rDNA contig was concatenated to the main chromosome arms displaying a telomeric sequence on one end and the 45S rDNA copies on the opposite end, creating a T2T chromosome with a single gap. The remaining unscaffolded and rDNA-devoid contigs, although accounting for 61 Mb in total, showed an extremely low average length (∼48 kb) as well as extremely low weighted coverage (∼14x), and thus they were also removed from the final assemblies (Supplementary Figure 4).

### Final Ajwa phased genome assembly

The Ajwa haplotype 1 assembly (Hap1) resulted in a final genome size of 673.1 Mb, with all telomeres present and a single gap located in the NOR region of Chr10. We found that the haplotype 2 assembly (Hap2) was of a very similar size at 670.1 Mb, with two gaps in the NOR region, an additional gap on Chr02, and one missing telomere on Chr03 (Figure 1a, Supplementary Figure 2). Based on these results, we consider the Ajwa reference genome to be complete and hereafter, refer to the Ajwa Hap1 assembly as the Ajwa reference genome (i.e. the primary assembly).

### A New Date Palm Chromosome Structure

Unexpectedly, a comparison of our new Ajwa reference genome with the first released Khalas PDK50^2^ genome revealed not only a smaller genome size, but more importantly, apparent major discrepancies at the chromosome level. Full details are provided in the Supplementary Note. In brief, we found that linkage groups (LGs) 4 and 16 in Khalas correspond to a single chromosome in Ajwa (Chr01), whereas LG11 in Khalas corresponds to two distinct, gap-free, and telomere-to-telomere (T2T) chromosomes in Ajwa (Chr09 and Chr18). Furthermore, all Khalas LGs appeared to be mosaics, composed of multiple Ajwa chromosomes. Multiple lines of evidence support the Ajwa chromosomal structure. We therefore suspected that the previous chromosome level assemblies needed curation. First, we previously demonstrated consistency with optical maps and Hi-C contact map patterns (Supplementary Figure 3). Second, as will be discussed in more detail in the following sections, we generated 19 additional high-quality date palm genomes (Figure 2a), all validated by Hi-C and Bionano optical maps, which display a chromosome architecture identical to that of Ajwa (Supplementary Figures 7 and 8). In addition, comparisons with other Aceraceae genomes, including American (*Elaeis oleifera* (Kunth) Cortés)^25,26^ and African (*Elaeis guineensis* Jacq.)^26,27^ oil palms, and *Cocos nucifera* L. (coconut palm)^27^, despite having different chromosome numbers, support the Ajwa chromosome structures for Chr01, Chr09, and Chr18 (Figure 4). To provide molecular evidence, we performed FISH experiments using probes designed on Chr01 to demonstrate that it represents a single continuous molecule. Likewise, cytological evidence for Chr09 and Chr18 confirmed that they represent two separate chromosomes. The structural discrepancies observed in the Khalas assembly can be attributed to the use of less advanced sequencing technologies and genetic map-based scaffolding strategies employed at the time. Indeed, the alignment of contigs from Khalas scaffolds onto the Ajwa genome supports the Ajwa chromosomal configuration as well and lacks the mosaic features. Similar results were obtained when comparing Ajwa and the Barhee BC4 genome (data not shown), which was generated using Khalas as a reference guide.

**Fig. 2.**
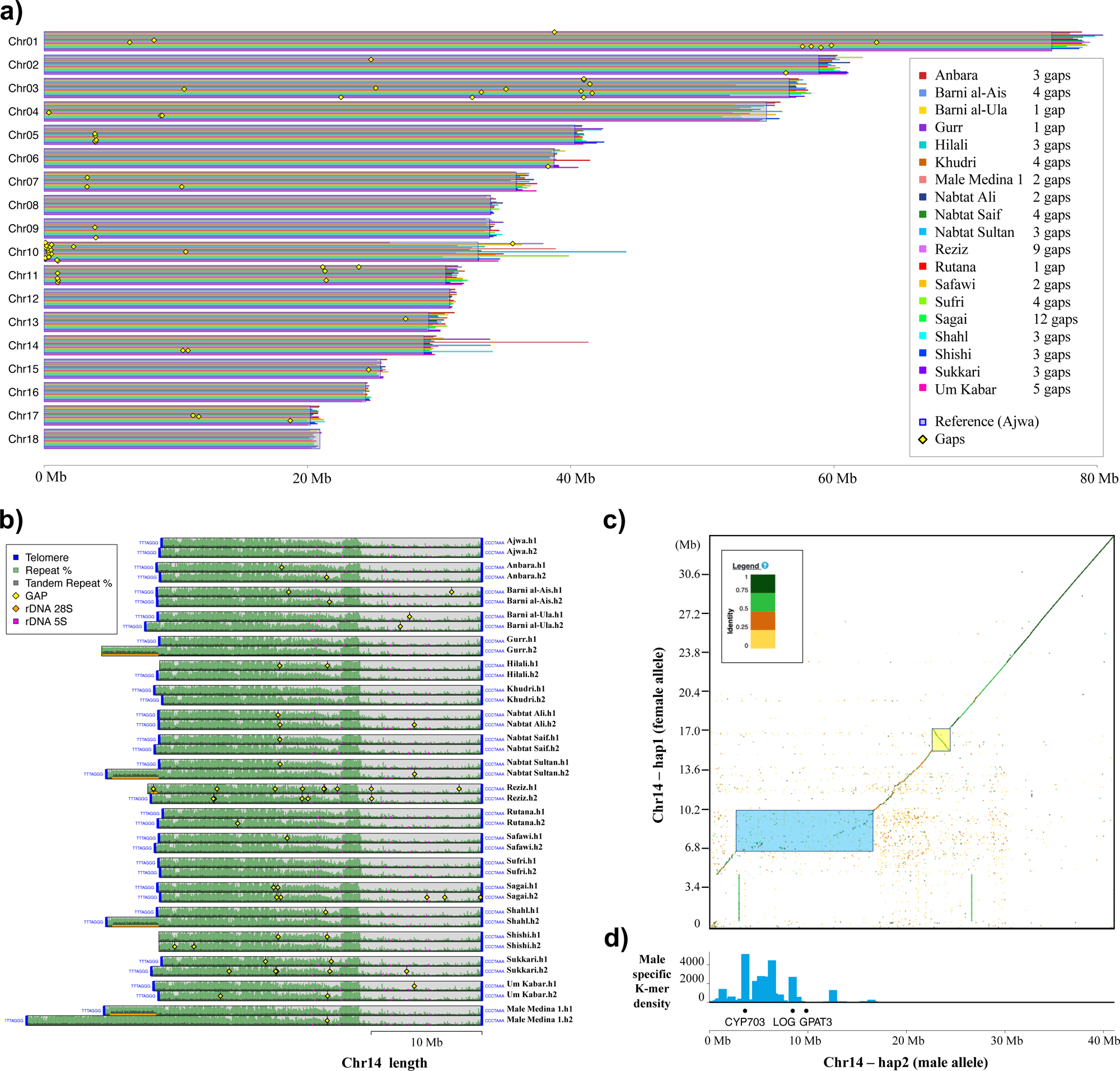
a) Scaffold length of 19 palm genomes created RagTag scaffold using Ajwa as a reference, manually curated and visualized with GS-viewer scaffold. b) Total repeat and tandem repeat profile of the all sex chromosomes (Chr14) of all 20 date palm genomes, shown as percentage in 100 Kb windows with the locations of telomeres, 5S rDNA and 45S rDNA, generated with GS-viewer repeat-profile. c) Dotplot of the Male date palm sex chromosomes created with D-Genies^48^, displaying the differential region in male and female chromosomes (yellow) and the male specific inversion (blue). d) Male specific K-mer counts per 100 Kb window (Hong *et al.,* in preparation) and male associated gene’s position from Torres *et al.*^36^.

Due to these ambiguities, and to unite the date palm community, we redefined the date palm chromosome numbering based on the average chromosome length observed in Ajwa and the other 19 new genome assemblies (Supplementary Figure 5), which are shown in Table 1.

### Repeat Profile

A custom repeat library for Ajwa was generated using EDTA v1.9.1^13^, and the genome was subsequently masked with RepeatMasker^28^. We found that 52.38% of the Ajwa genome consisted of repetitive elements, with the majority represented by LTR retrotransposons (42.61%), mainly from the Copia (24.34%) and Gypsy (12.35%) families. Copia elements were particularly enriched in the central regions of chromosomes. In contrast, Gypsy elements showed a more uniform distribution throughout the genome. Notably, Chr11 displayed a localized enrichment of unclassified LTR retrotransposons not observed elsewhere in the genome. DNA transposons accounted for 7.51% of the genome, with the DTM/Mutator superfamily being the most prevalent component, mostly co-localizing with the ribosomal repeats in the NOR region of Chr10 (Supplementary Figure 6).

As previously mentioned, no centromeric/pericentromeric tandem repeat patterns were identified. As noted, Copia elements were enriched in the central regions of each chromosome, but not in a centromeric type distribution as seen with other centromeric TEs ^29,30^.Thus, we do not expect that these elements correspond to the functional centromeres. We also employed the CentiER pipeline^31^ to identify potential centromeric repeat makers, but no candidate centromere sequences were found. These analyses suggest that Date palm centromeres may possess a novel sequence structure and will require a date palm CENH3 specific antibody for further investigation.

### Assembly of 19 Additional Date Palm Genomes (18 females and 1 male)

Following the assembly, phasing, and validation of the Ajwa genome, we generated genome assemblies for 19 additional date palm accessions, including one male genet and 18 commercially important female cultivars grown from Saudi Arabia: Anbara, Barni al-Ais, Barni al-Ula, Gurr, Hilali, Khudri, Nabtat Ali, Nabtat Saif, Nabtat Sultan, Reziz, Rutana, Safawi, Sufri, Sagai, Shahl, Shishi, Sukkari, and Um Kabar (Table 2). The male sample was collected from the same location as Ajwa; the al-Dabta farm in Medina, KSA. The Barni accession from al-Ula, which had already been short-read sequenced by Gros-Balthazard & Battesti *et al.*^32^ under the ID 00067, was sampled in a private farm garden in the al-Ula oasis (project al-Ula DPA). The remaining 17 female cultivars were obtained from the National Center for Palms and Dates (NCPD) in al-Ahsa, KSA. As with Ajwa, we generated high-quality assemblies using PacBio HiFi long reads together with HiC reads, assembled with Hifiasm v0.24.00, and further validated each assembly with Bionano optical maps. Sequencing and assembly metrics were consistent with those of Ajwa, although some assemblies showed slightly higher fragmentation (Table2, Supplementary Tables 1-2). K-mer analyses revealed consistent profiles across all accessions, supporting a diploid genome feature with high heterozygosity similar to that of Ajwa and included in 0.5-0.7% (Supplementary Figure 1). Scaffolding of the 19 assemblies was carried out using RagTag Scaffold ^33,34^, employing the Ajwa genome as a reference guide. A custom tool, GS-viewer (https://github.com/mirkocelii/GS-viewer), was run to generate chromosome-specific plots for the validation of contig alignment on the reference and the removal of misplaced and duplicated contigs. In line with the Ajwa strategy, the multiple 45S rDNA repeat contigs on chromosome 10 were not concatenated; the chromosome was closed only with a telomere-bearing contig obtained from either the primary assembly or a phased haplotype, when available. Chloroplast and mitochondrial sequences were aligned to the Ajwa organellar genomes with RagTag Scaffold and manually curated with Genome Puzzle Master^35^ when required.

**Table 2.**
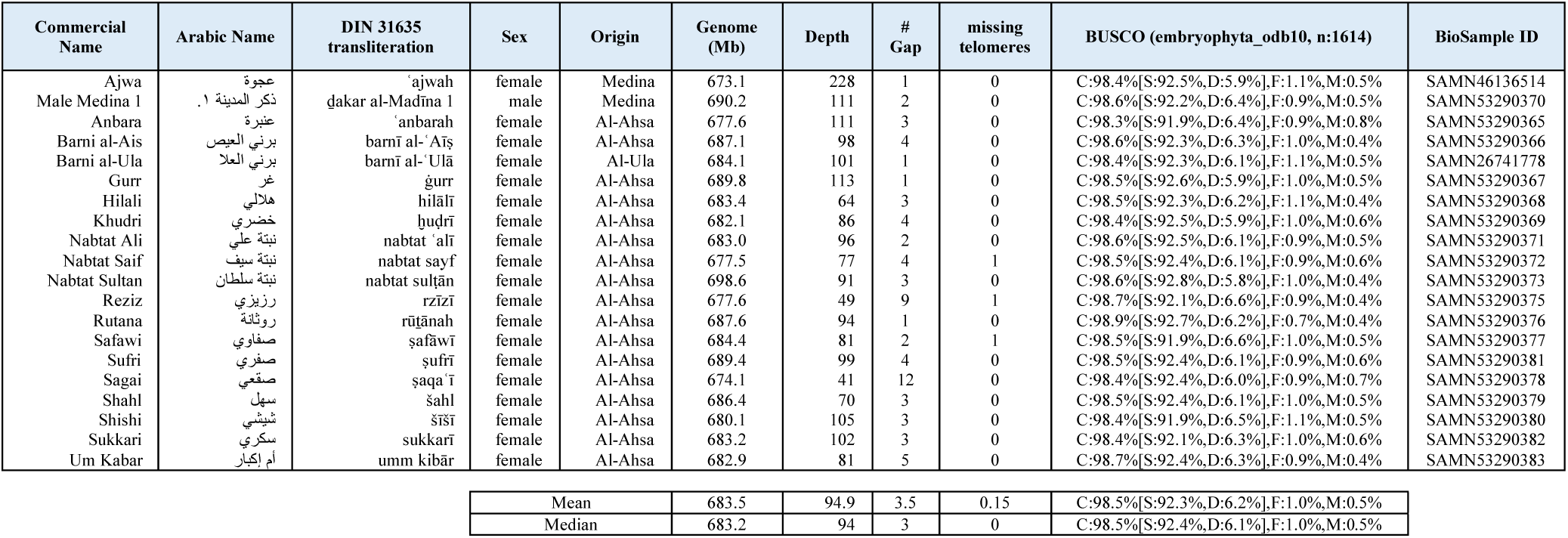
Date palm genomes used in this study, with their Arabic cultivar names, transliterations according to the DIN 31635 system, sex, sample origin, assembly statistics and BioSample accessions. *The male individual has no commercial cultivar name and is referred to by its sample ID. Gaps in organelle assemblies are not included in the gap counts.

All 19 genomes, including the male, displayed strong collinearity with the Ajwa genome assembly, exhibited a low number of assembly gaps (ranging from 1 to 12), a consistent repeat profile (Supplementary Figure 7) and are telomere-to-telomere complete, except for three assemblies in which a single telomere was absent. Three female genomes, and the female haplotype of the male genome contained a secondary 45S locus of ∼5.7 Mb in length on chromosome X (Figure 2a-b). Each assembly was also validated with a high-quality hybrid scaffold with Bionano optical maps (Supplementary Figure 8).

### 45S rDNA TE-Related Spacer

Another notable feature of the 45S rDNA array was the presence of a TE-derived spacer (#<u>TE_00001883)</u> sequence between each 45S rDNA unit, that had sequence homology to *Mutator*-like DNA transposable element terminal inverted repeats (Figure 3b). These repeated spacer sequences represent one of the most abundant repeats in the genome by copy number (Supplementary Figure 4) and were found exclusively on NOR located on Chr10, in all rDNA-rich contigs, and occasionally on Chr04, while being completely absent from the rest of the genome assembly (Figure 3a). (Supplementary Figure 9). The exclusive localization of this sequence within NORs was confirmed by FISH (Figure 3d) using a cloned (2.1-TOPO TA vector (Invitrogen)) PCR amplified 1,789 bp fragment of the repeat (forward: 5’-GTCGTGACAGTTTTCGGTGG-3’; reverse: 5’-GTACTTTGGAACGCTTGGGG-3’). This clone was then labeled with either digoxigenin-11-dUTP or biotin-16-dUTP using a standard nick translation reaction and processed as described for the rDNA 45S and telomere probes. Although the presence of transposon-related rDNA spacers has been described before^39,40^, their existence and the extent of these repeats in the date palm genome could only be revealed here due to the application of the most recent long-read sequencing technologies and assembly tools.

**Fig. 3.**
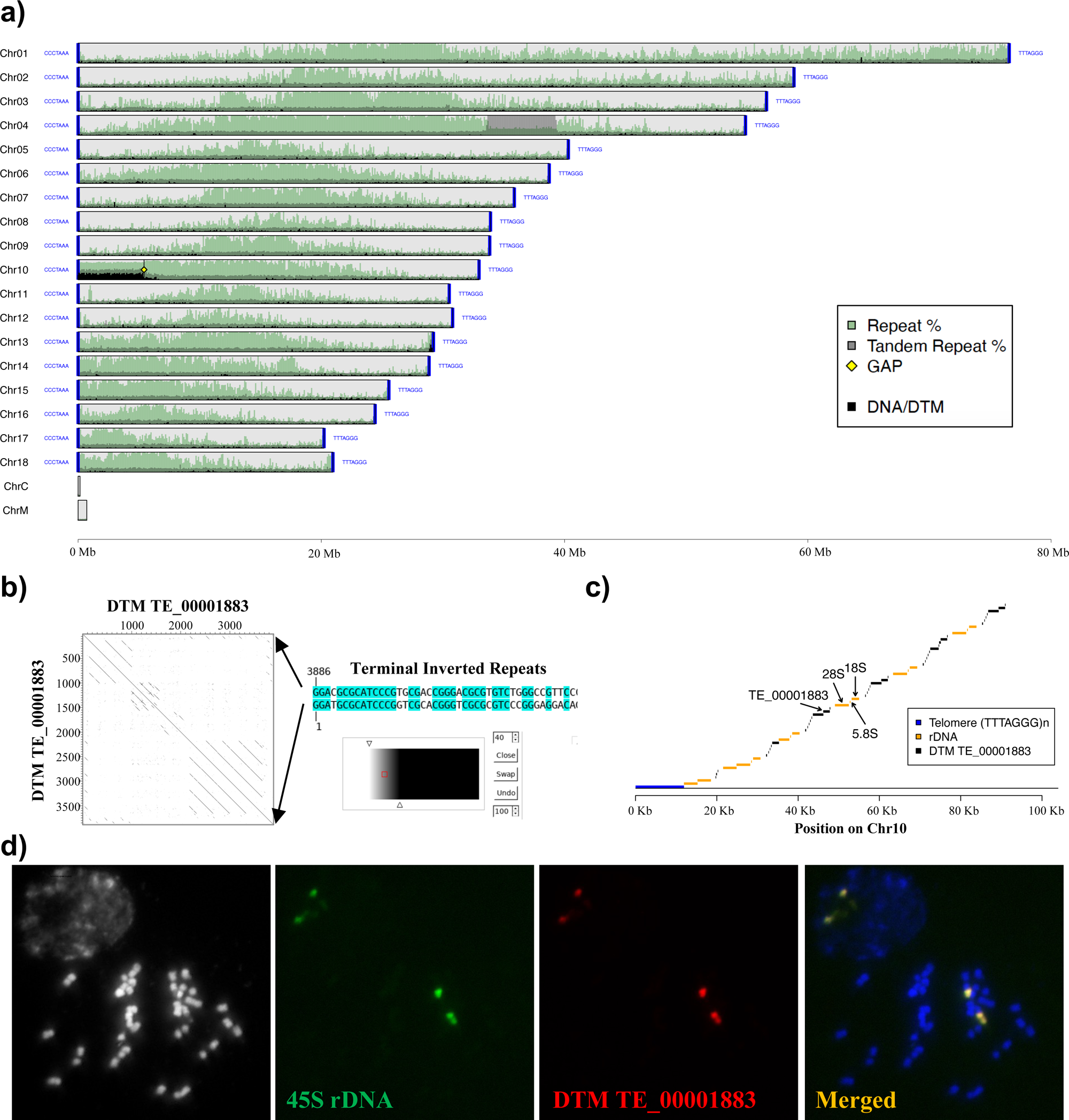
a) Background: Total repeat and tandem repeat content in 100Kb windows; Foreground DNA/DTM TE_00001883 counts in 100Kb windows, generated with GS-viewer repeat-profile. b) Dotplot of the putative non-autonomous DNA Mutator-like element created with Dotter^52^. c) Representation of the concatenation pattern of the 3 ribosomal RNAs and the spacer sequence including the TE_00001883 repeat. d) Validation of exclusive co-localization of rDNA 45S loci and TE_00001883 on the same chromosome by FISH using rDNA 45S and TE_00001883 specific probes.

### Genome Size and Completeness

The 20 date palm assemblies exhibited highly consistent genome sizes ranging between 673-689 Mb (mean = 683.5 Mb; median = 683.2 Mb) with most of the observed variation attributable to differences in the length of the 45S rDNA regions on chromosomes 04, 10, and in a few sex chromosomes carrying the additional 45S rDNA locus (Figure 2a, Table 2, Supplementary Figure 8). These genome sizes are significantly smaller than those previously reported for Khalas (854 Mb)^2^ and Barhee BC4 (772 Mb)^4^ and the flow cytometry-based estimate of 843 Mb^21^.To assess genome completeness, we ran BUSCO^41^ using the embryophyta_odb10 database. All 20 genomes showed high and consistent completeness, with at least >98.3% of complete BUSCO genes and an average of ∼6% duplication. These results highlight a more accurate estimate of the date palm genome size in the range of 670–700 Mb. The larger genome sizes reported in earlier assemblies likely reflect higher levels of duplication (e.g. Barhee BC4 ∼12%, Khalas ∼9.4% of duplicated BUSCOs) and the presence of substantial unplaced sequence fractions (13% and 46%, respectively) (Supplementary Figure 10).

### Gene Prediction and Annotation

Gene prediction was performed across all 20 date palm assemblies using the OmicsBox 3.4^42^ wrapper, that integrated AUGUSTUS 3.5.0^43^, HTSlib 1.19.1^44^ BCFtools 1.19 ^45^, Samtools 1.19.2 ^46^, and HDF 1.12.2^47^ tools with *Oryza sativa* L. as the reference species in the *ab initio* mode. For the Ajwa genome specifically, we integrated both RNA-seq data and Iso-Seq datasets from roots, leaves, and flowers. RNA-seq data were merged, and a subsample of 30M reads was used for gene prediction. The predicted gene count was highly consistent across all 20 genomes, ranging from 32,770 to 33,996 genes. As expected, the Ajwa annotation, which incorporated transcriptomic evidence, resulted in a higher gene count of 41,264. Similar to genome size variation, gene number differences were associated with the variable length of the 45S rDNA region on Chr10, and occasionally on sex chromosome Chr14 (Supplementary Figure 11).

### Date Palm Sex Determining Region

Previous studies have shown an XY-like sex determination system in date palm, and the identification of three candidate male-specific genes on Khalas LG12 (homologous to Ajwa Chr14): *CYP703* and *GPAT3*, which are absent in the female haplotype, and *LOG3*, which is thought to have originated through duplication and translocation from an autosome. Additionally, a chromosomal inversion on the male chromosome has been suggested to suppress recombination^36,37^. To validate the results from these studies we used blastn^38^ to search for the three male-specific genes across the 19 female and one male genome assemblies. As expected, *CYP703* and *GPAT3* could be found on the male haplotype of Chr14 and were absent in all female Chr14 assemblies. Of interest, *LOG3* was found on autosome Chr16 in both male and female genomes; however, a second *LOG3* locus (with 86% sequence identity) could be found on the male haplotype of Chr14 (Figure 2d, Supplementary Table 4). Together these results are consistent with previous studies identifying the putative candidate genes for sex determination in date palm.

The sex-determining region (SDR) in date palm was estimated by previous studies to range in size from 6 to 13 Mb^3,4^. We mapped male-specific k-mers (Hong *et al.,* in preparation, BioProject ID PRJNA1278712) that uniquely align to non-homologous regions of the male haplotype of Chr14, which spanned approximately ∼14.7 Mb of Chr14 and encompassed all three male-specific genes (i.e. *CYP703*, *GPAT3*, and *LOG3)*. The non-homologous region of the female haplotype was much smaller in length at ∼4.7 Mb (Figures 2c–d). Lastly, we identified a 1.6 Mb male-specific inversion on Chr14, that was absent in all female accessions sequenced, consistent with Torres *et al.*^36,37^ (Figure 2c, Figure 5).

Of interest, in addition to the three male specific genes previously identified, and confirmed here, gene prediction analysis (see details below) of the 14.7 Mb SDR revealed the presence of 45 additional genes; 30 harbor the previously described male-specific k-mers (21 with Gene Ontology [GO] annotation), 19 lack a detectable counterpart on the female chromosome 14 (9 with GO annotation), and 4 satisfy both criteria (1 with GO annotation). This expanded gene content may shed new light on the sex determination mechanism(s) in date palm in future studies (Supplementary Figure 12).

### Comparison with Other Palm Genomes

As previously mentioned, we observed that the genomes of other palm species, including the American and African oil palms (*Elaeis oleifera*^25^, BioProject ID PRJNA183707; *Elaeis guineensis*^26^, BioProject ID PRJNA192219) and coconut palm (*Cocos nucifera*^27^, BioProject ID PRJNA413280), are structurally consistent with the Ajwa genome assembly, despite differing in chromosome number. We used the online tool D-Genies^48^ to generate both whole-genome and chromosome-specific alignments. Ajwa Chr01 (homologous to Khalas LG04 and LG16) corresponded to a single chromosome in both oil and coconut palms. Similarly, Chr09 and Chr18 (both homologous to Khalas LG11) correspond to two separate chromosomes in all three species. Interestingly, Chr18 appears fused with another chromosome in both oil and coconut palms, specifically with the sex chromosome Chr14 in *C. nucifera*, and with Chr10 in *E. guineensis* (Figure 4). While the synteny of these three date palm chromosomes with *E. guineensis* had already been reported^3^, the lower resolution of their data did not allow for the identification of the distinct chromosome architecture observed here for date palm.

**Fig. 4.**
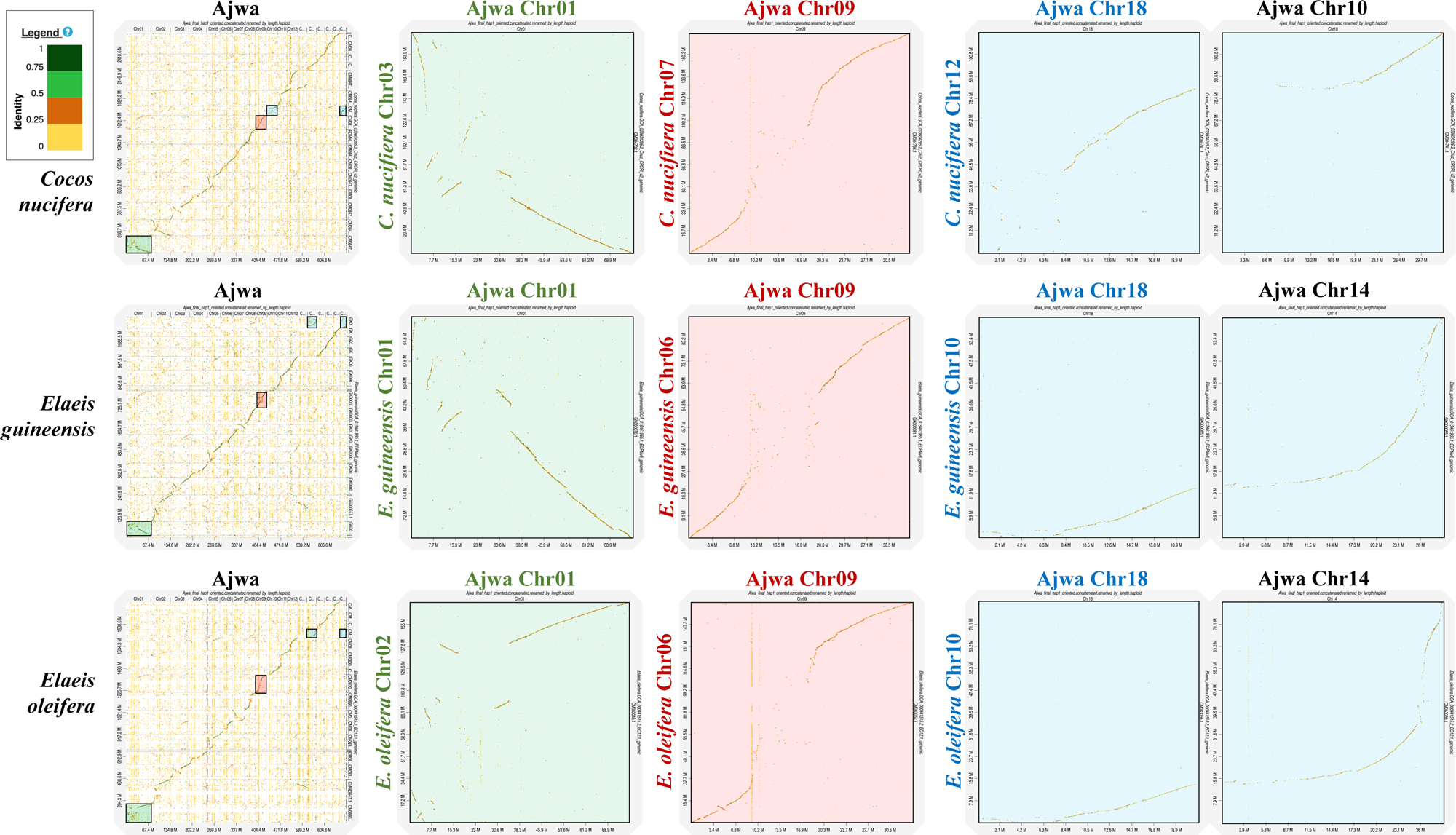
Dotplot of Ajwa genome compared *to Cocos nucifera*^27^, *Elaeis guineensis*^26^ *and Elaeis oleifera*^25^ and of Ajwa Chr01, Chr09 and Chr18 with their homologous in the other 3 palm genomes, created with D-Genies^48^.

### Gene Synteny

To investigate and visualize syntenic relationships across the 19 Saudi date palm varieties and the male genet, we generated a synteny map including all 20 date palm assemblies and the outgroup species *E. guineensis* (Figure 5). Gene coordinates and protein sequences were used in GENESPACE v.1.3.1^49^, which embeds OrthoFinder v.2.5.4^50^ and MCScanX_h^51^ using default settings, to define orthogroups. The resulting riparian plot shows strong collinearity within the *Phoenix dactylifera* genomes. Interestingly, a number of large-scale inversions were observed, particularly in putative centromeric and pericentromeric regions on Chr06, Chr01, Chr05, Chr10, and Chr03. Sex chromosome (Chr14) showed no inversions in all female genomes, but a distinct inversion was observed in the male haplotype, consistent with Torres *et al.*^36,37^. Interestingly, no large-scale translocations involving annotated genes were detected.

**Fig. 5.**
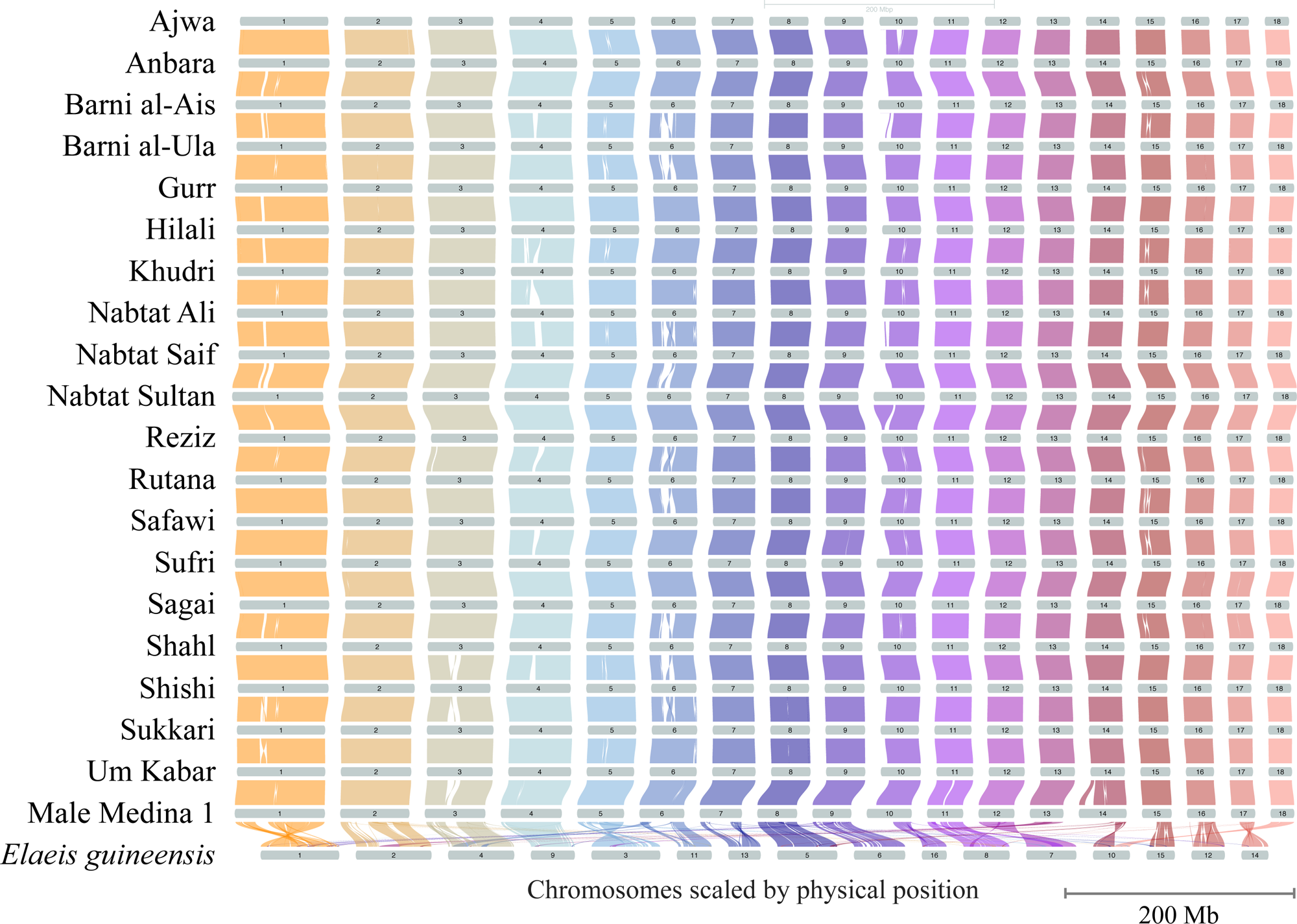
Riparian plot showing macro-syntenic regions and large-scale structural rearrangements (large duplications and translocations) across the chromosomes of 20 date palm genomes and the outgroup species *Elaeis guineensis*^26^.

## Conclusion

Together, the Ajwa genome and 19 additional telomere-to-telomere (T2T) assemblies resolve chromosome-scale structure, refine the estimated genome size to 680–700 Mb, and reveal a gap-free ∼14.7 Mb male sex-determining region (SDR). To unify the date palm genomics community, we propose that the Ajwa assembly with additional male haplotype sequence for the SDR described here be considered as the reference assemblies for future date palm research.

## Data Records

All data have been deposited in GenBank (https://www.ncbi.nlm.nih.gov/) under the BioProject ID PRJNA1207559. Biosamples IDs of each date palm accession are indicated in Table 2. Barni al-Ula data were deposited under the existing biosample ID SAMN26741778 generated by Gros-Balthazard & Battesti *et al.*^32^ Bionano Hybrid Scaffolds, Gene predictions and the Ajwa repeat library were deposited on Zenovo [10.5281/zenodo.17684092]

Male-specific k-mers and all associated sequencing data were generated as part of a companion study on early male individual identification, which is in preparation for publication, whose raw data are deposited under NCBI BioProject PRJNA127871

## Technical Validation

### DNA and RNA Sample Quality

Nucleic acid concentrations were quantified with a Qubit fluorometry Quantification system (Invitrogen, USA), and a NanoDrop spectrophotometer (Thermo Fisher Scientific, USA). To evaluate DNA integrity and restriction accessibility, samples were digested with EcoRI and HindIII restriction enzymes, followed by gel electrophoresis. DNA fragment size distributions were then estimated using a Femto Pulse System (Agilent Technologies, USA).

### Pacbio Libraries

Pacbio Library concentration was measured using the Qubit Fluorometer High-Sensitivity Kit (Invitrogen), and fragment size distribution was assessed on the Femto Pulse System (Agilent Technologies, USA)

### Illumina Libraries

Illumina Libraries were quantified by qPCR using the KAPA Library Quantification Kit for Illumina Libraries (KapaBiosystems, Wilmington, MA, USA), and library profiles were evaluated with an Agilent 2100 Bioanalyzer (Agilent Technologies, Santa Clara, CA, USA).

### Gene Space Completeness

Benchmarking Universal Single-Copy Orthologs (BUSCO3.0) was run using the embryophyta_odb10 database to assess the gene completeness of all 20 genomes (Table2, Supplementary Figure 11).

### Assembly Accuracy

Bionano optical maps and HiC contact maps were used to validate all 20 genome assemblies. (Supplementary Figure 3, Supplementary Figure 7).

### Chromosome Structure

Florescence *in situ* Hybridization (FISH) was performed to validate chromosome number, telomeric sequences, NOR. NOR-related spacer, and the structure of Chr01, Chr08 and Chr18 (Figure 1a, Figure 3d, Supplementary Note Figure 2c)

## Code Availability

All tools used in this study are listed below. When parameters are not specified, the tool was run using default settings.

GS-viewer code is available on GitHub at https://github.com/mirkocelii/GS-viewer:

– Repeat Profiles were generated with GS-viewer repeat-profile
– Scaffolds created with RagTag were visualized with GS-viewer scaffold.

K-mer analysis: jellyfish count -C -s 5G -m 51; genomescope2 -k 52 -p 2 PacBio Read analyses: seqkit stats -aT

Hi-C and RNAseq reads trimming: Trimmomatic

Genome Assembly: (Ajwa Phased): hifiasm/0.19.0 --n-weight 6 --n-perturb 10000 -s 0.40; (Other 19 primary assemblies): hifiasm/0.24.0 --n-weight 6 -s 0.40

TE annotation: edta/1.9.7 -species others

Contamination removal: Kraken2: extract_kraken_reads.py --taxid 33154 2 2157 --include-children

Organelle identifications: OATK

Tandem repeat annotation: Tandem Repeat Finder: trf 2 3 5 80 10 30 2000 -l 6 -dat -h Repeat Masking: Repeat Masker: RepeatMasker -qq -no_is

Scaffolding:

– RagTag: ragtag.py scaffold -f 70000 -a 0.005 -u -w --remove-small
– D-Genies
– Bionano Solve: hybridScaffold.pl -B 2 -N 2 -f
– YAHS

TE sequence analysis: Dotter

Gene annotations: Omicsbox: reference *Oryza sativa*, default parameters

Riparian Plot: GENESPACE v.1.3.1, default settings; plot_riparian()

## Acknowledgements

This research was supported by the King Abdullah University of Science & Technology through the Center for Desert Agriculture (CDA) under the Center Project Funding (CPF): *Fast Fit Palms*; KAUST baseline funding to JP and RW, the National Center for Palms and Dates (NCPD) in al-Aḥsa, KSA; The al-Dabta farm in Medina, KSA; and the French Agency for al-Ula Development (AFALULA) with its Saudi partner, the Royal Commission for al-Ula (RCU), through a grant awarded to V.B. and M.G-B. (*project al-Ula DPA: Ethnographic, genetic, and morphometric analyses of the date palm agrobiodiversity in al-Ula oasis*). Legally, the Royal Commission for al-Ula is the owner and provider of the material of the Barni variety from al-Ula.

## Ethics Statement

This work was approved by the King Abdullah University of Science and Technology (KAUST), the National Center for Palms and Dates (NCPD). All methods used in this study were carried out following approved guidelines.

## Author contributions

I.B., J.P., and R.A.W. designed and conceived the research.

M.N.R. and U.T. prepared libraries for PacBio and Illumina sequencing.

N.A., Y.Z., M.N., and R.S. worked on the initial Ajwa genome draft.

V.L. generated and assembled Bionano optical maps.

K.F. generated the phased Ajwa genome.

M.C. generated curated genomes of the other 18 date palms varieties and the male accession.

M.C. ran Ajwa TE annotation and gene predictions for all 20 date palm genomes.

M.C. run the comparative analyses of Barhee BC4, Khalas PDK50 and other palm genomes.

M.C. created GS-viewer.

D.K. generated the FISH images.

A.F. performed gene syntenic analysis.

Y.H. performed the male k-mer analysis.

M.C. submitted data.

M.G.B., T.C., M.P. and J.A.M. advised, consulted and consented on chromosome renaming.

I.B., M.G.B. and V.B. contributed to the sampling.

M.C., J.P., and R.A.W. wrote and edited the paper.

J.P., and R.A.W. contributed equally as senior authors.

All authors read and approved the final manuscript.

