## Supplementary Figures for "Foundational genomic resources for date palm: A gap-free, telomere-to-telomere phased assembly of Ajwa and 19 high-quality genome assemblies of *Phoenix dactylifera*"

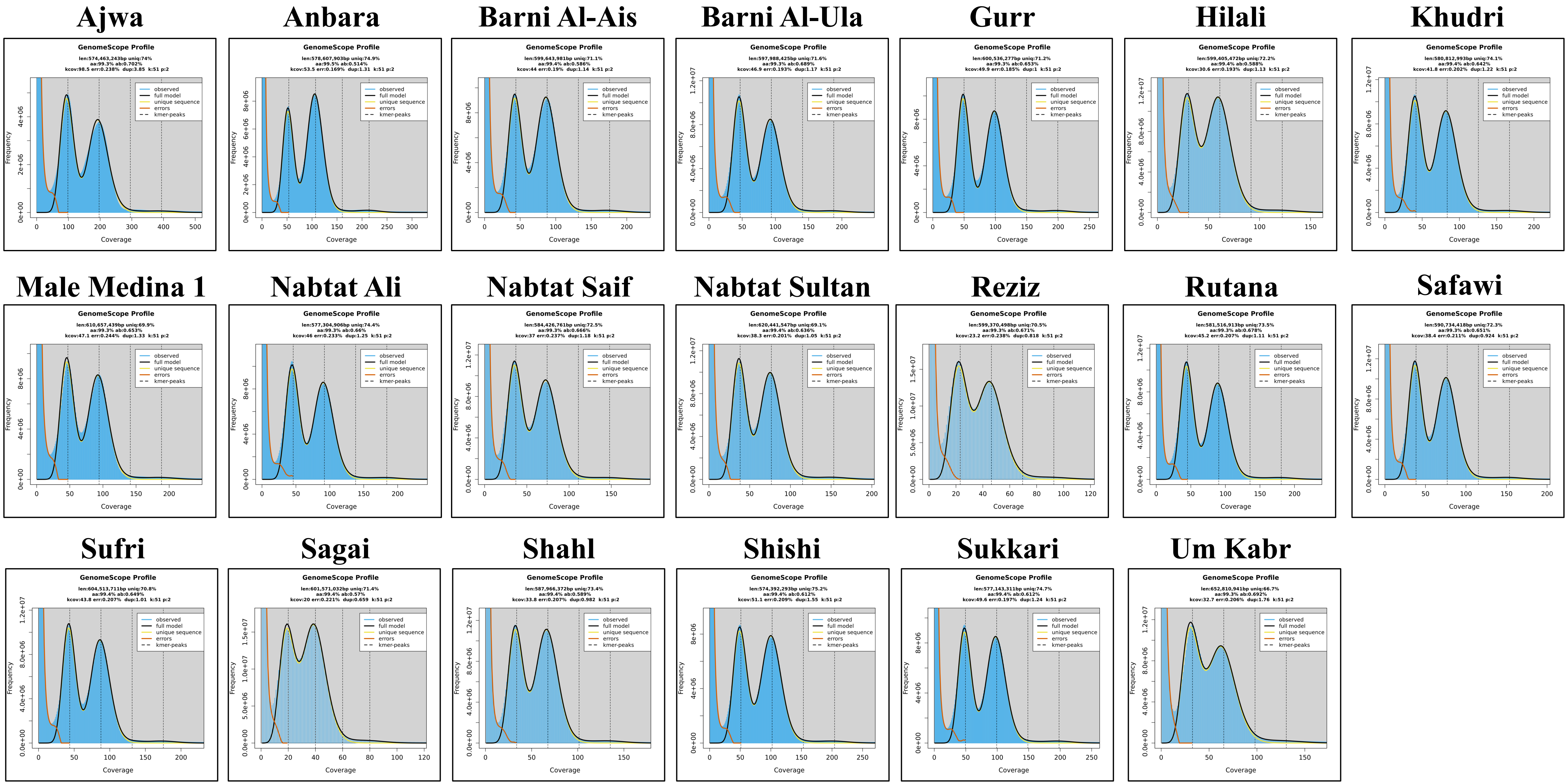


**Supplementary Figure 1.** Kmer profile of all 20 Date Palm accessions generated with Jellyfish v2.3.0[14] and visualized GenomeScope v2.0[15] using k-mer length of 51 bp

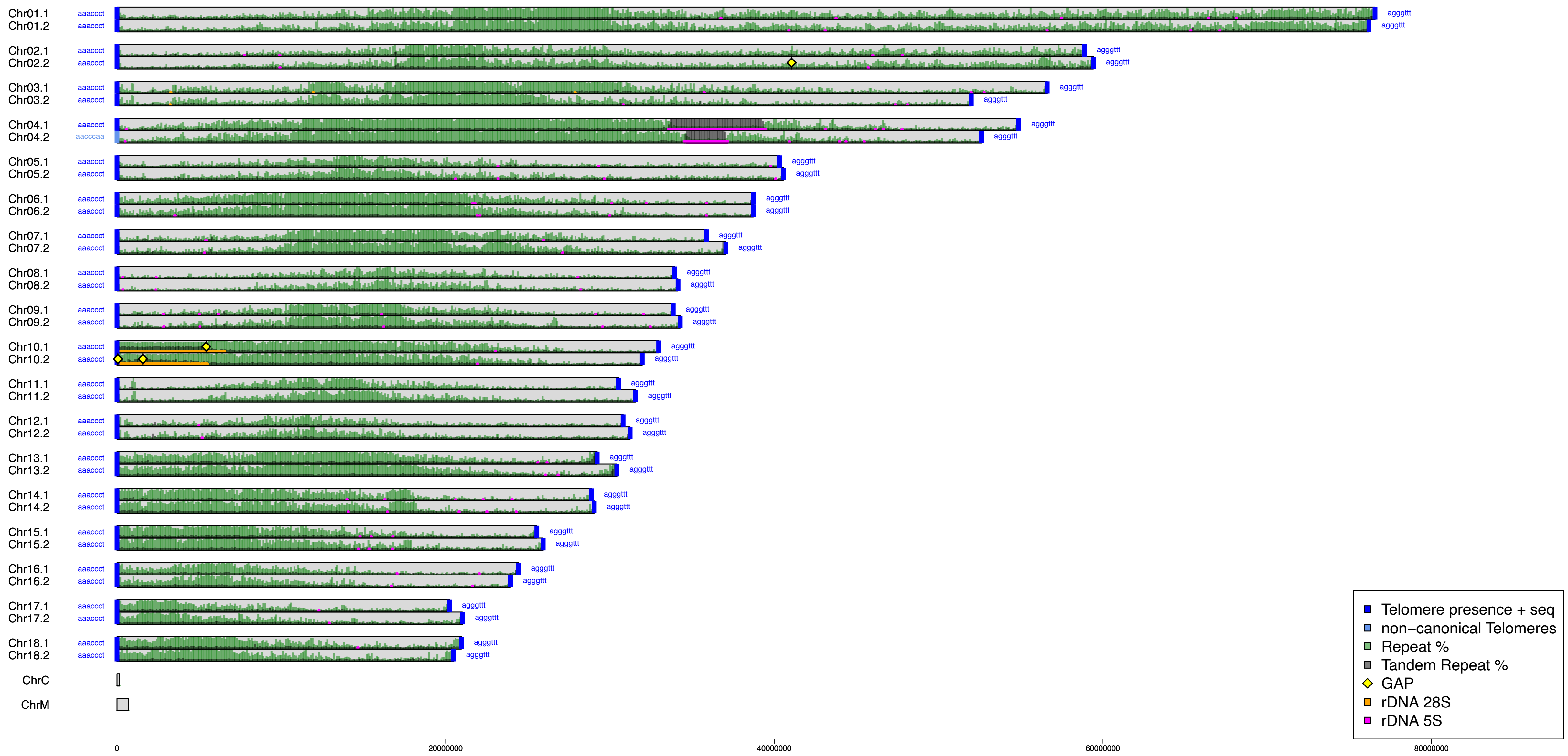


**Supplementary Figure 2.** Total repeat and tandem repeat profile shown as percentage in 100 Kb windows with presence of telomere, rDNA 5S and rDNA 45S locations in the Ajwa diploid genome realized with GS-viewer.R repeat-profile

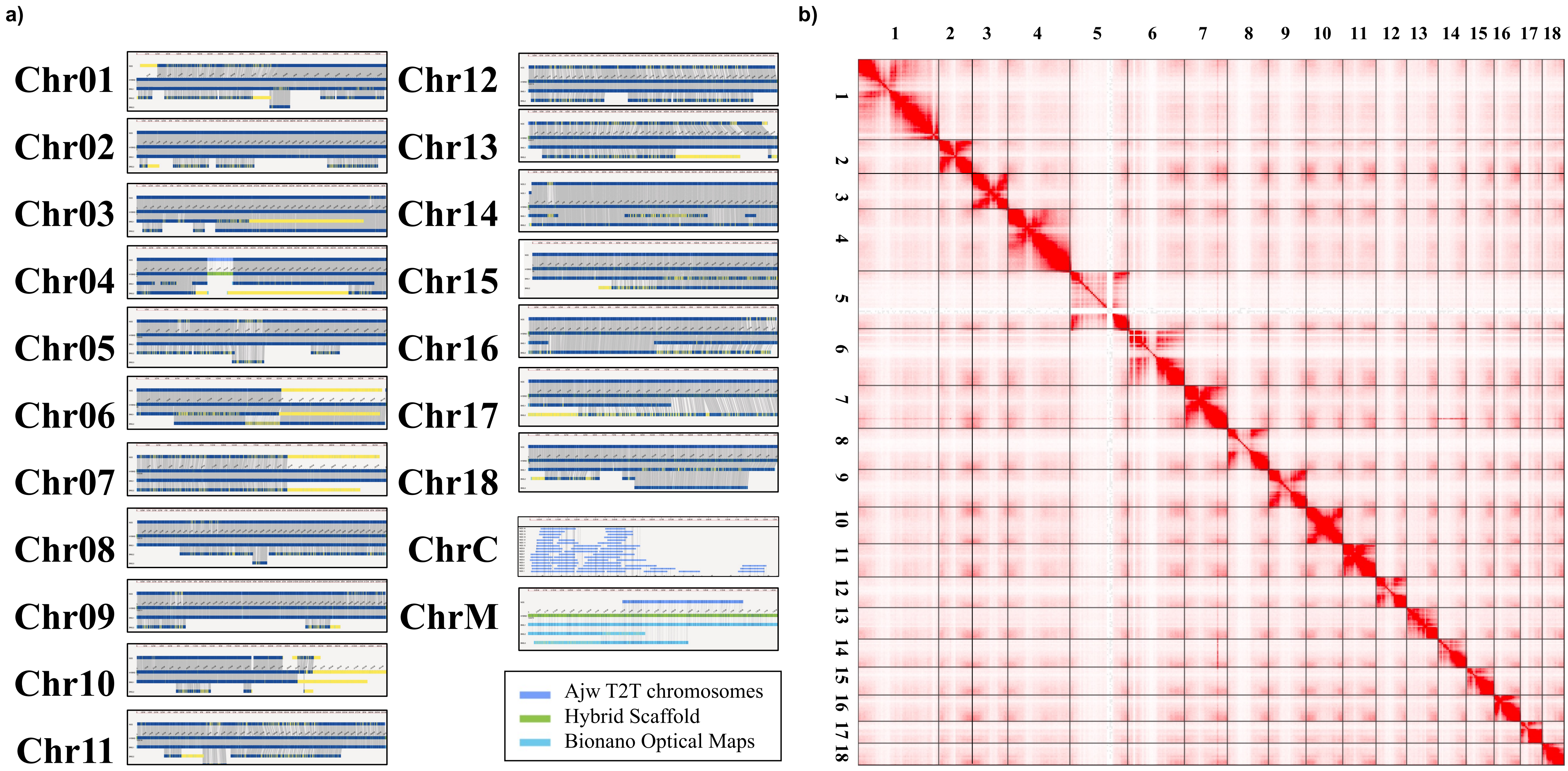


**Supplementary Figure 3.** a) Snapshot from Bionano Access viewer of 20 hybrid scaffold of Ajwa hap1 matching with the 18 chromosomes and the organellar genomes. Input used for the 18 Chromosomes was the Ajwa hap1 genome, input for the organelles was the unscaffolded and unfiltered assembly. b) Hi-C contact map of Ajwa hap1 generated with YAHS[24] and visualized with Juicebox

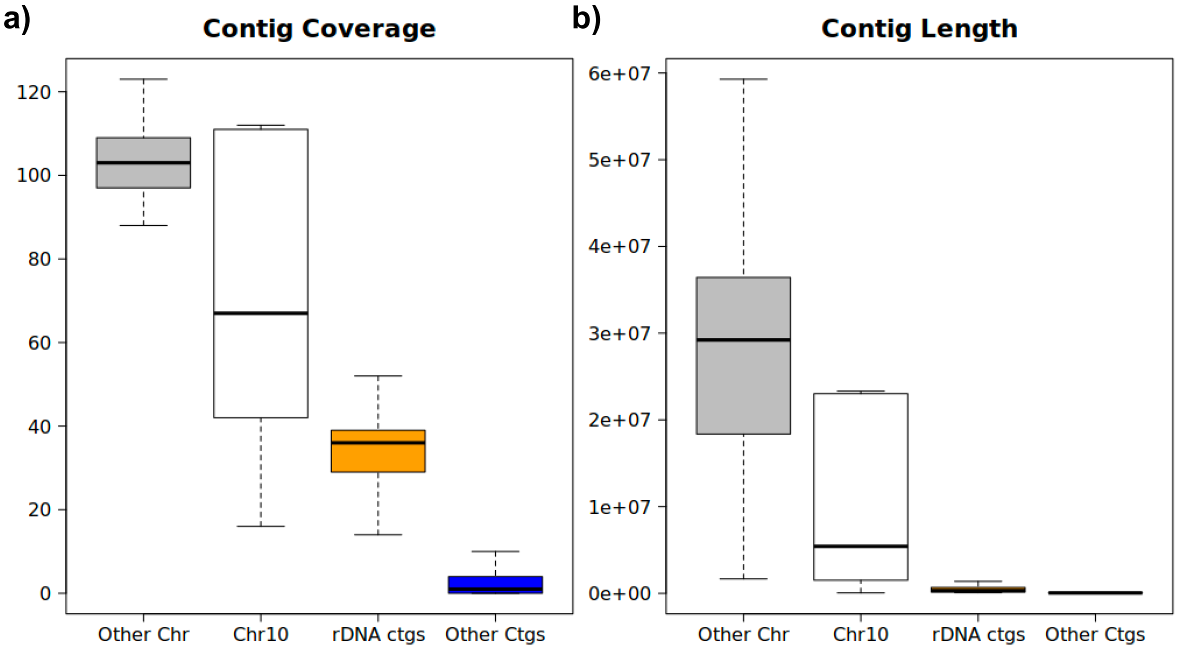


c)

| | Contigs | | | | |
|---|---|---|---|---|---|
| | **All Chr.** | **Chr10** | **Other Chr.** | **rDNA unplaced** | **other unplaced** |
| **Size in Mb** | **1333.06** | **53.34** | **1279.7** | **75.92** | **61.32** |
| **Weighted coverage** | **107.64** | **103.07** | **107.83** | **45.8** | **14.15** |

**Other Chr / rDNA cov. ratio = 2.25**

**Estimated rDNA length per haplotype (Mb) = 16.87**

**Supplementary Figure 4.** a) Distribution of contig coverages extracted from Hifiasm's hap1 and hap2 *.gfa files, divided in 4 groups: unplaced rDNA-rich contigs, placed contigs on NOR carrying Chr10, placed contigs on other chromosomes, unplaced contigs devoid of rDNA copies. b) Distribution of contig length of the same 4 groups in a). c) Estimation of rDNA total length by weighting the coverage of rDNA contigs on the chromosome-placed contigs.

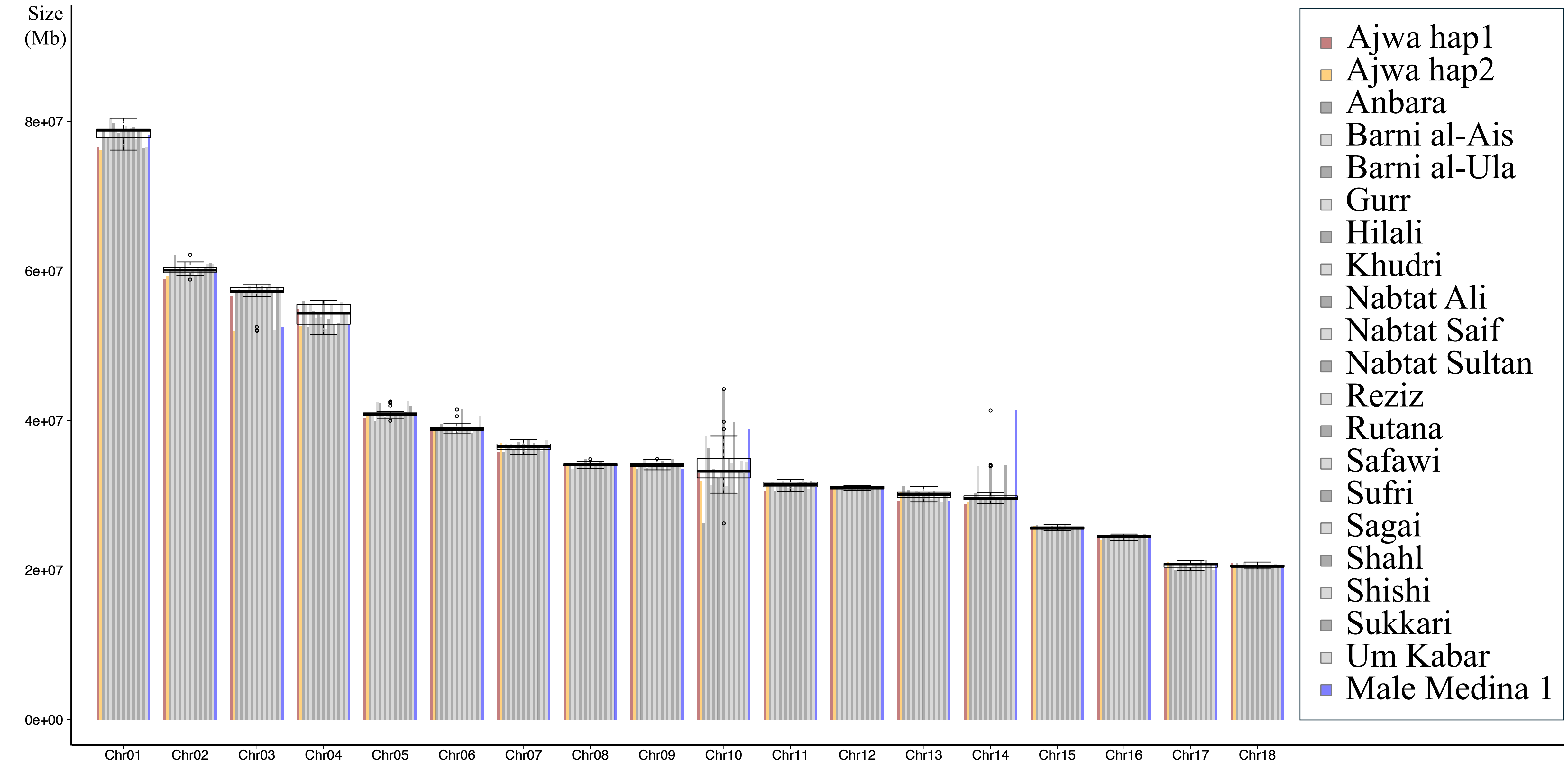


**Supplementary Figure 5.** Chromosome length of Ajwa phased haplotypes and the other 19 date palm primary genomes. Each bar represents individual values; Boxplot are representing median, quartiles maximum and minimum values for each chromosome

**a)**

| Elements | number of elements | length occupied | percentage of sequece |
|---|---|---|---|
| Retroelements: | 275216 | 283732190 bp | 42.1 % |
| SINEs: | 0 | 0 bp | 0 % |
| Penelope: | 0 | 0 bp | 0 % |
| LINEs: | 0 | 0 bp | 0 % |
| CRE/SLACS | 0 | 0 bp | 0 % |
| L2/CR1/Rex | 0 | 0 bp | 0 % |
| R1/LOA/Jockey | 0 | 0 bp | 0 % |
| R2/R4/NeSL | 0 | 0 bp | 0 % |
| RTE/Bov-B | 0 | 0 bp | 0 % |
| L1/CIN4 | 0 | 0 bp | 0 % |
| LTR elements: | 275216 | 283732190 bp | 42.1 % |
| BEL/Pao | 0 | 0 bp | 0 % |
| Ty1/Copia | 127607 | 164725967 bp | 24.44 % |
| Gypsy/DIRS1 | 78537 | 83241722 bp | 12.35 % |
| Retroviral | 0 | 0 bp | 0 % |
| DNA transposons: | 165534 | 50630837 bp | 7.51 % |
| hobo-Activator | 0 | 0 bp | 0 % |
| Tc1-IS630-Pogo | 0 | 0 bp | 0 % |
| En-Spm | 0 | 0 bp | 0 % |
| MULE-MuDR | 0 | 0 bp | 0 % |
| PiggyBac | 0 | 0 bp | 0 % |
| Tourist/Harbinger | 0 | 0 bp | 0 % |
| Other | 0 | 0 bp | 0 % |
| Rolling-circles: | 0 | 0 bp | 0 % |
| Unclassified: | 51344 | 10099889 bp | 1.5 % |
| Total interspersed repeats: | | 344462916 bp | 51.11 % |
| Small RNA: | 0 | 0 bp | 0 % |
| Satellites: | 0 | 0 bp | 0 % |
| Simple repeats: | 136870 | 7075061 bp | 1.05 % |
| Low complexity: | 27159 | 1527091 bp | 0.23 % |

| | | |
|---|---|---|
| **Total length:** | **673990002 bp** | |
| **Sequences:** | **20** | |
| **Total base masked:** | **353065068 bp** | **52.38 %** |
| **GC level:** | | **40.61 %** |

**b)**

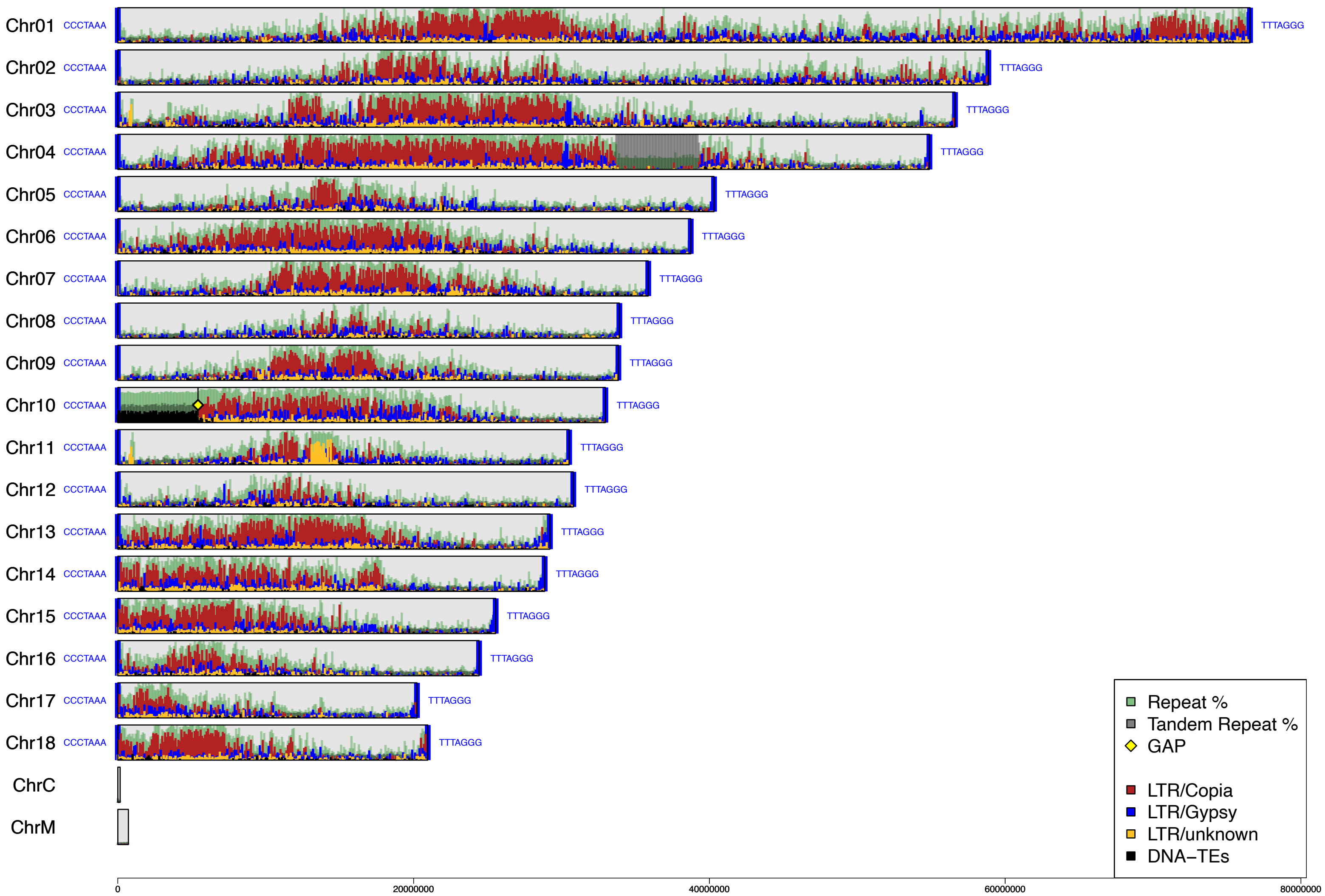


**Supplementary Figure 6.** a) Repeat Masker statistics on Ajwa genome b) Repeat profile of Ajwa genome, in background: total repeat and tandem repeat content in 100Kb windows; in foreground: LTR Copia, LTR Gypsy, LTR Unknown and DNA-TE profiles in 100Kb windows, generated with GS-viewer repeat-profile.

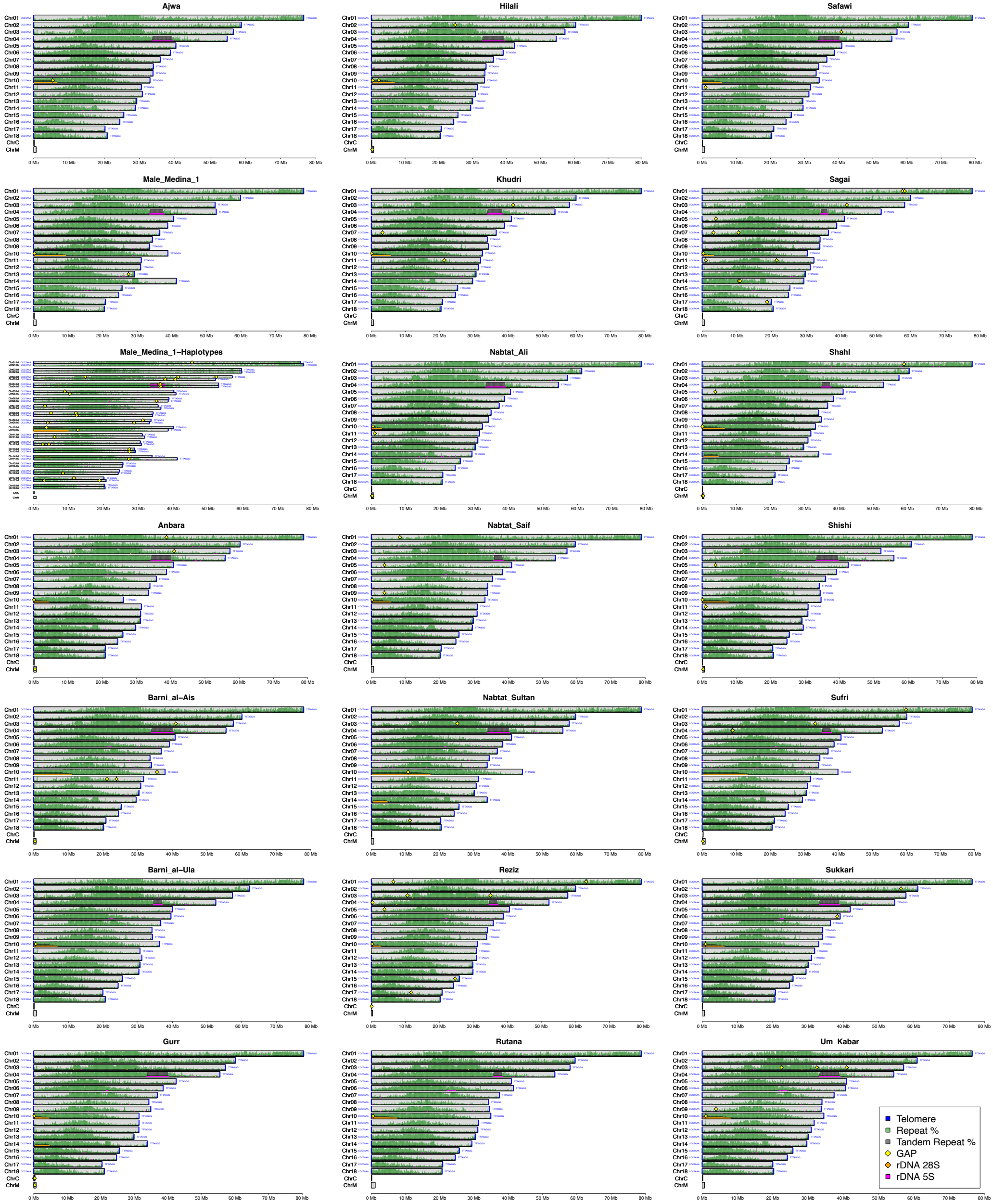


**Supplementary Figure 7.** Total repeat and tandem repeat content in 100Kb windows with telomeres, gaps, rDNA 5S and rDNA 45S locations of 20 date palm genomes and of the 2 haplotype of the Male Medina 1 genome

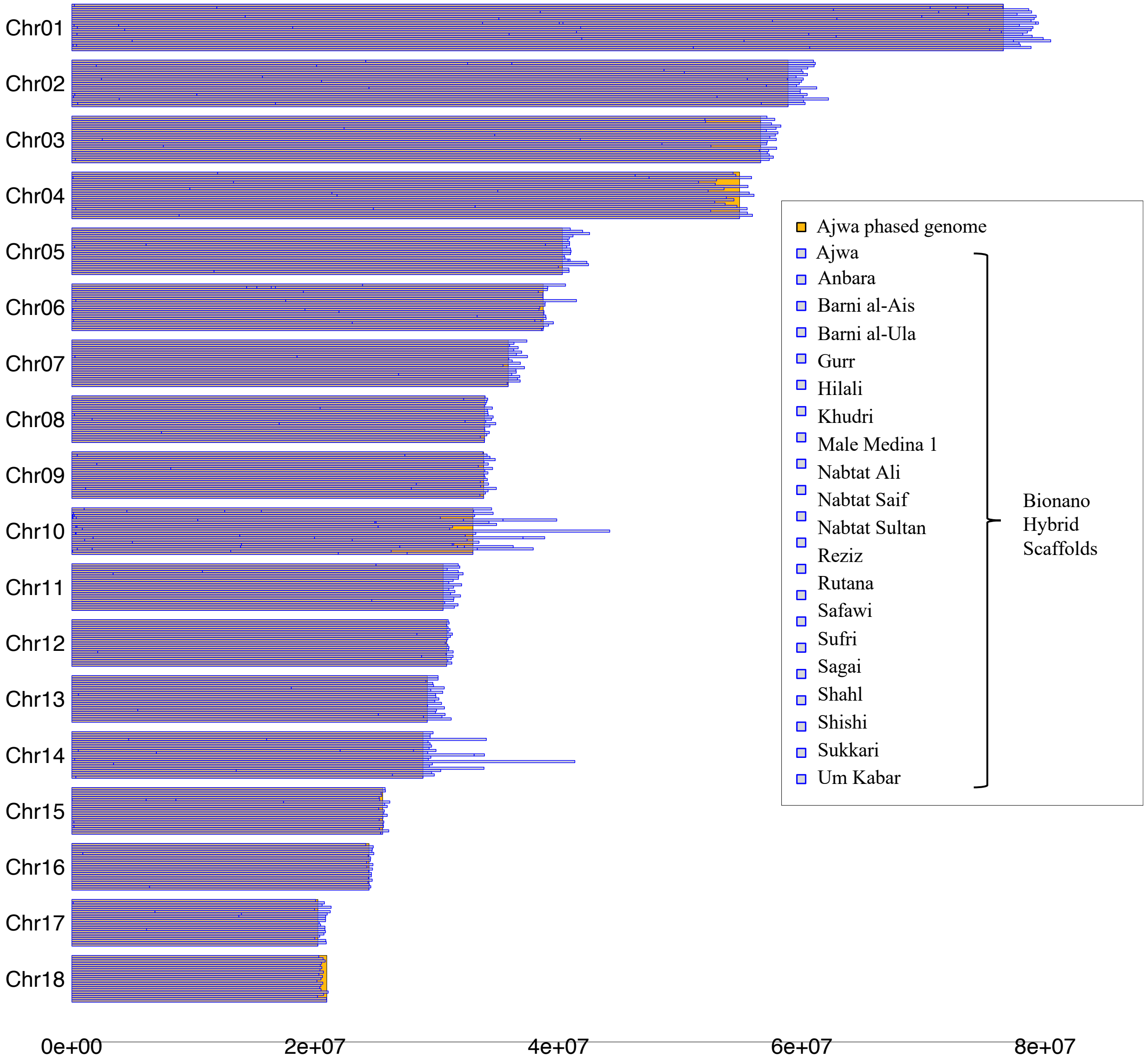


**Supplementary Figure 8.** Representation of the 20 bionano hybrid scaffold based on the AGP files generated by Bionano solve (blue) compared to the Ajwa genome (yellow)

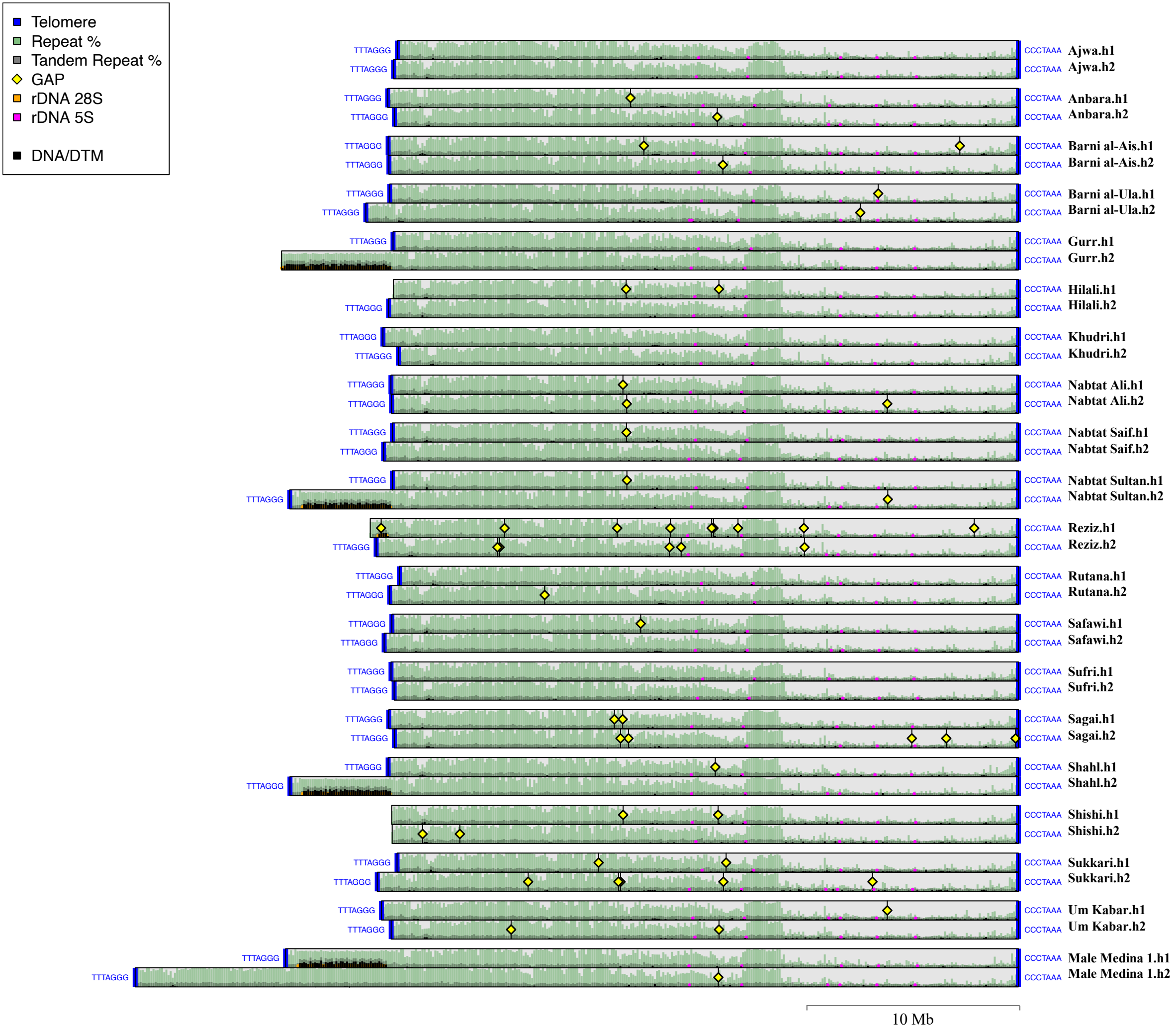


**Supplementary Figure 9.** Repeat profile of sex chromosomes of the 20 date palm genomes. Background: Total repeat and tandem repeat content in 100Kb windows in all date palm sex chromosomes; Foreground DNA/DTM TE_00001883 counts in 100Kb windows, generated with GS-viewer repeat-profile.

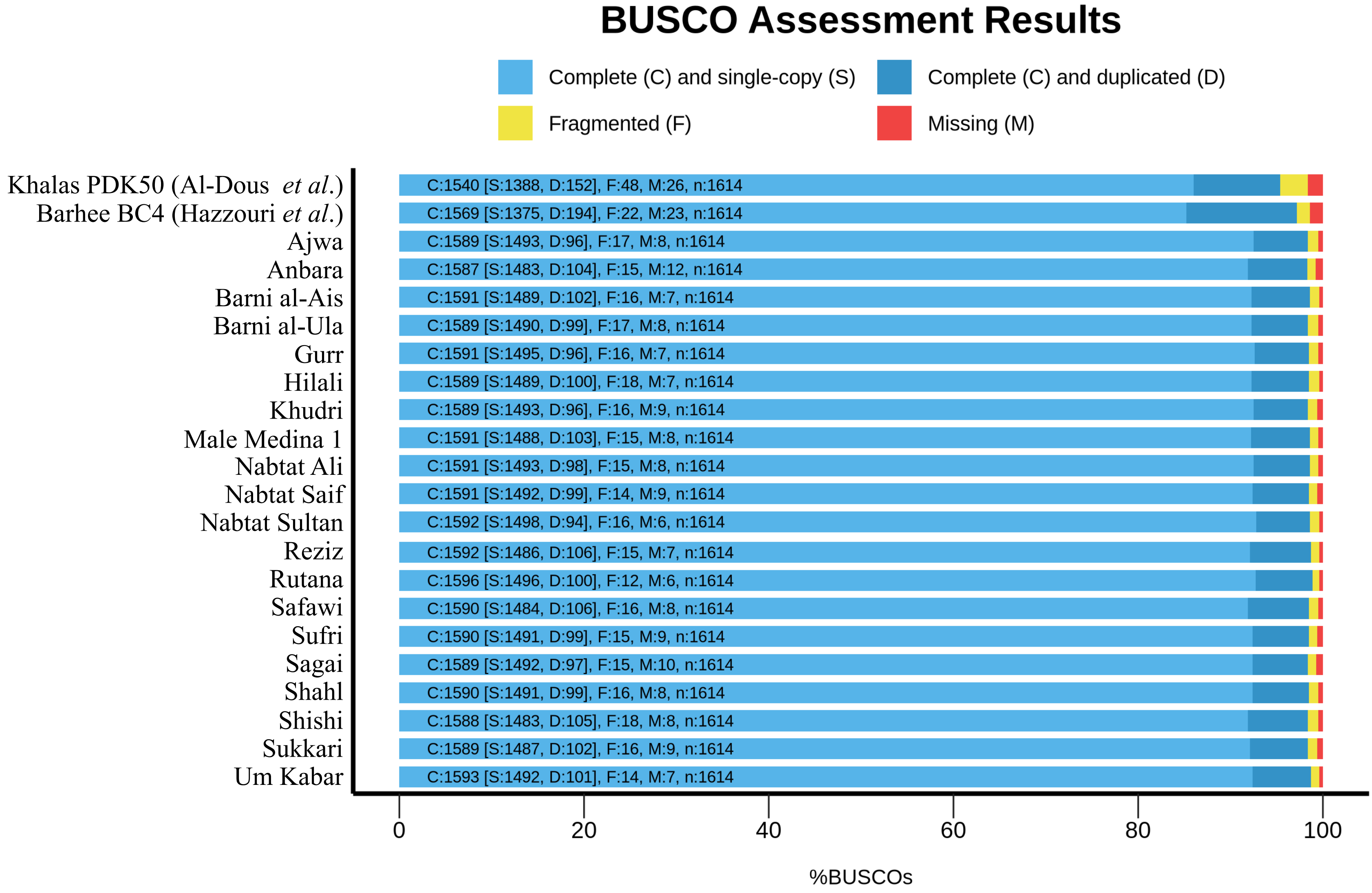


**Supplementary Figure 10.** Genome completeness evaluation with BUSCO using the embriophytsa_obd10 database of 20 date palm genomes compared to the previously released Khalas PDK50 (Al-Dous *et al.,* 2011 updated in 2020) and Barhee BC4 (Hazzouri *et al.,* 2019)

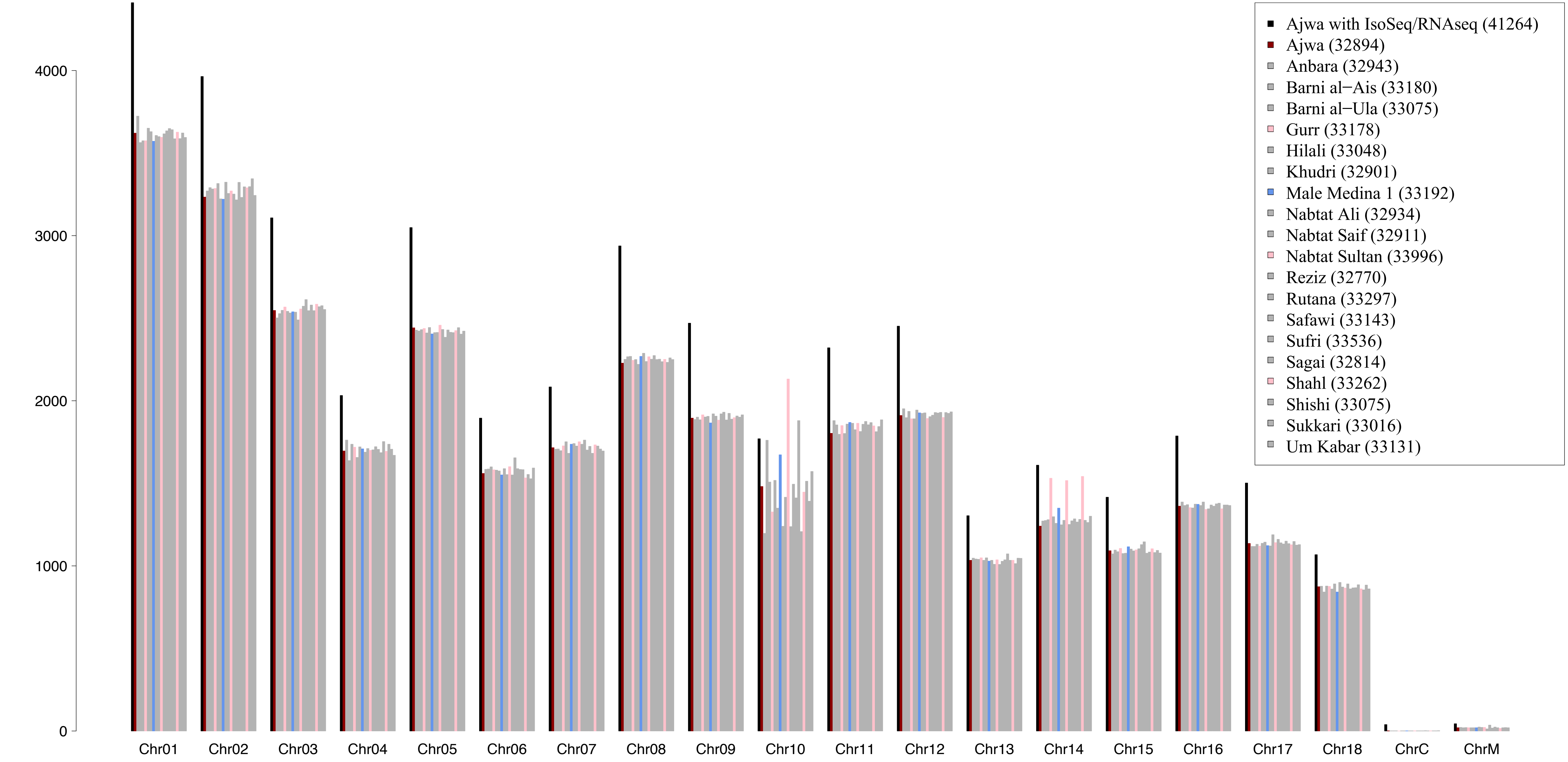


**Supplementary Figure 11.** Predicted Gene number by Omicsbox for Ajwa (including and not including RNAseq an IsoSeq reads) and the other 19 genomes (predicted genes only). Black: Ajwa including RNA-seq and ISOseq; Brown: Ajwa with predicted genes only, Blue: Male genome; Pink: female genomes with second NOR localized in the Sex Chr10; Gray: all other female genomes.

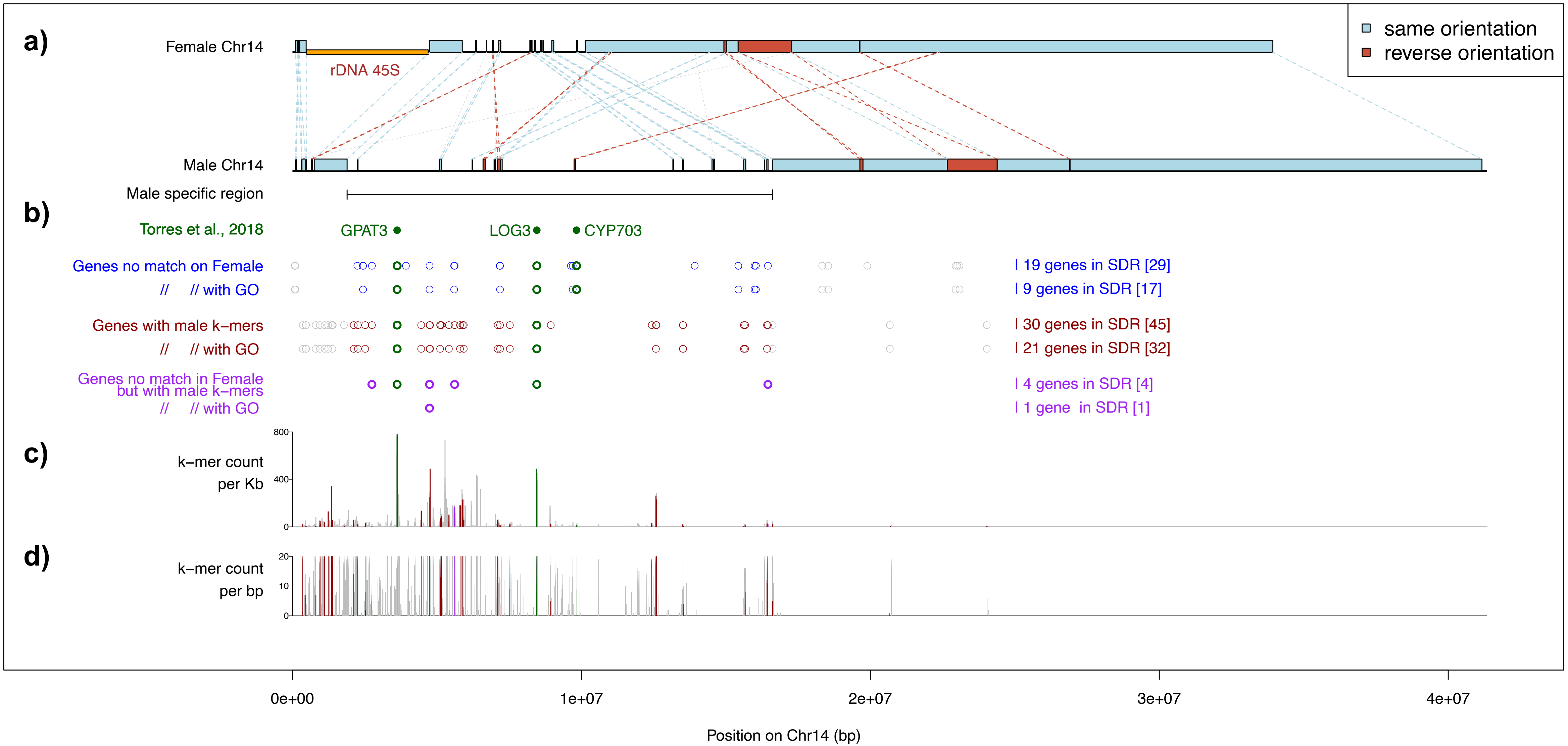


**Supplementary Figure 12.** a) Representation of the Sex-Determining Region (SDR) in the male and female haplotypes of the Male Medina 1 date palm. Homologous regions (light blue and red boxes) were identified using RagTag scaffolding. The male SDR is defined as the region that shows no detectable homology with the female chromosome. b) Genes located within the male SDR. Green: SDR genes from Torres *et al.*: *GPAT3* (Gene ID: LOC120108105), *LOG3* (Gene ID: LOC103701078) and *CYP703* (Gene ID: LOC120107104), identified by BLASTN. Blue: male genes with no homologues on the female chromosome. Dark red: male genes overlapping male-specific k-mers (20 bp) from Hong *et al.* (in preparation). Purple: male genes that both lack homologues on the female chromosome and overlap male-specific k-mers. Grey background: genes that lack homologues in the female haplotype but are located outside the putative SDR. c) Density of male-specific k-mers per 1 kb window, using the same colour scheme as in b) to indicate the corresponding gene categories. d) K-mer counts per base pair, using the same colour scheme as in (b).

| Accessions | file names | Platform | | format | type | num_seqs | sum_len | min_len | avg_len | max_len | Q1 | Q2 | Q3 | sum_gap | N50 | Q20(%) | Q30(%) | GC(%) |
|---|---|---|---|---|---|---|---|---|---|---|---|---|---|---|---|---|---|---|
| Ajwa | m64041_210217_190046.hifi_reads.fastq.gz | AGI | Sequel II | FASTQ | DNA | 952217 | 19027141418 | 45 | 19981.9 | 51075 | 17969 | 19863 | 22232 | 0 | 20463 | 97.69 | 94.54 | 41.3 |
| Ajwa | m64041_210219_012206.hifi_reads.fastq.gz | AGI | Sequel II | FASTQ | DNA | 879117 | 17481582210 | 45 | 19885.4 | 50925 | 17937 | 19855 | 22233 | 0 | 20479 | 97.79 | 94.79 | 41.33 |
| Ajwa | m64041_210223_063711.hifi_reads.fastq.gz | AGI | Sequel II | FASTQ | DNA | 967685 | 18524714094 | 46 | 19143.3 | 52851 | 17381 | 19328 | 21566 | 0 | 19940 | 97.64 | 94.47 | 40.81 |
| Ajwa | m64068_221115_134005.hifi_reads.fastq.gz | KAUST BCL | Sequel II | FASTQ | DNA | 1852132 | 34958580567 | 130 | 18874.8 | 50600 | 14457 | 18017 | 22374 | 0 | 19939 | 98.51 | 96.54 | 40.74 |
| Ajwa | m64468e_221115_133940.hifi_reads.fastq.gz | KAUST BCL | Sequel IIe | FASTQ | DNA | 1615289 | 30758217669 | 124 | 19041.9 | 50438 | 14647 | 18204 | 22533 | 0 | 20098 | 98.58 | 96.64 | 40.81 |
| Ajwa | m64468e_221116_230519.hifi_reads.fastq.gz | KAUST BCL | Sequel IIe | FASTQ | DNA | 1867446 | 35287215237 | 139 | 18896 | 50357 | 14501 | 18049 | 22381 | 0 | 19962 | 98.55 | 96.56 | 40.69 |
| Ajwa Isoseq | AjwaFlower.m64468e_220708_073046.hifi_reads.bc2003--bc2003.bam.fastq | KAUST BCL | Sequel II | FASTQ | RNA | 819099 | 2071221409 | 121 | 2528.7 | 10988 | 1947 | 2360 | 2913 | 0 | 2619 | 99.29 | 98.31 | 45.35 |
| Ajwa Isoseq | AjwaLeaf.m64468e_220708_073046.hifi_reads.bc2004--bc2004.bam.fastq | KAUST BCL | Sequel II | FASTQ | RNA | 972893 | 2689304380 | 99 | 2764.2 | 16068 | 2115 | 2588 | 3203 | 0 | 2856 | 99.24 | 98.2 | 43.59 |
| Ajwa Isoseq | AjwaRoot.m64468e_220708_073046.hifi_reads.bc2005--bc2005.bam.fastq | KAUST BCL | Sequel II | FASTQ | RNA | 1161874 | 2881566775 | 96 | 2480.1 | 14308 | 1881 | 2290 | 2872 | 0 | 2580 | 99.29 | 98.32 | 45.42 |
| Anbara | Anbara_rep1.ccs.fastq.gz | KAUST BCL | Sequel II | FASTQ | DNA | 2258813 | 38143124728 | 132 | 16886.4 | 50104 | 13821 | 16247 | 19361 | 0 | 17301 | 98.62 | 96.68 | 40.79 |
| Anbara | Anbara_rep2.ccs.fastq.gz | KAUST BCL | Sequel II | FASTQ | DNA | 2266070 | 37335501378 | 133 | 16475.9 | 50303 | 13499 | 15817 | 18849 | 0 | 16836 | 98.67 | 96.83 | 40.78 |
| Barni al-Ais | m64468e_230129_144038.hifi_reads.fastq.gz | KAUST BCL | Sequel IIe | FASTQ | DNA | 1841372 | 32944412593 | 98 | 17891.2 | 49588 | 14476 | 17240 | 20717 | 0 | 18522 | 98.63 | 96.79 | 40.66 |
| Barni al-Ais | m64468e_230130_235428.hifi_reads.fastq.gz | KAUST BCL | Sequel IIe | FASTQ | DNA | 1942779 | 34294835014 | 106 | 17652.5 | 50431 | 14274 | 17018 | 20481 | 0 | 18311 | 98.62 | 96.78 | 40.5 |
| Barni al-Ula | m64068_221212_141456.hifi_reads.fastq.gz | KAUST BCL | Sequel II | FASTQ | DNA | 1999650 | 34026819691 | 88 | 17016.4 | 49265 | 13637 | 16399 | 19808 | 0 | 17695 | 98.4 | 96.2 | 40.9 |
| Barni al-Ula | m64468e_221212_141209.hifi_reads.fastq.gz | KAUST BCL | Sequel IIe | FASTQ | DNA | 1982840 | 34891790945 | 111 | 17596.9 | 49478 | 14094 | 17026 | 20564 | 0 | 18380 | 98.67 | 96.86 | 40.37 |
| Gurr | m64468e_230201_091606.hifi_reads.fastq.gz | KAUST BCL | Sequel IIe | FASTQ | DNA | 2170348 | 39181060666 | 114 | 18052.9 | 50465 | 14851 | 17512 | 20751 | 0 | 18607 | 0 | 0 | 41.06 |
| Gurr | m64468e_230202_183723.hifi_reads.fastq.gz | KAUST BCL | Sequel IIe | FASTQ | DNA | 2132359 | 38018349364 | 113 | 17829.2 | 49505 | 14670 | 17268 | 20464.5 | 0 | 18343 | 0 | 0 | 41.04 |
| Hilali | Halali_rep1.ccs.fastq.gz | KAUST BCL | Sequel II | FASTQ | DNA | 1285522 | 23456529201 | 86 | 18246.7 | 50655 | 14894 | 17584 | 21003 | 0 | 18769 | 98.61 | 96.74 | 40.61 |
| Hilali | Halali_rep2.ccs.fastq.gz | KAUST BCL | Sequel II | FASTQ | DNA | 1096370 | 19914727512 | 86 | 18164.2 | 50040 | 14828 | 17496 | 20897 | 0 | 18670 | 98.43 | 96.36 | 40.57 |
| Khudri | Khodri_rep1.ccs.fastq.gz | KAUST BCL | Sequel II | FASTQ | DNA | 1481009 | 24035801623 | 86 | 16229.3 | 49605 | 13551 | 15707 | 18481 | 0 | 16577 | 98.48 | 96.46 | 40.5 |
| Khudri | Khodri_rep2.ccs.fastq.gz | KAUST BCL | Sequel II | FASTQ | DNA | 2147502 | 34818326229 | 78 | 16213.4 | 50168 | 13541 | 15686 | 18452 | 0 | 16554 | 98.72 | 97.08 | 40.65 |
| Male_Medina_1 | m64468e_230404_121158.hifi_reads.fastq.gz | KAUST BCL | Sequel IIe | FASTQ | DNA | 1775670 | 36929644527 | 181 | 20797.6 | 50693 | 16662 | 20094 | 24248 | 0 | 21674 | 0 | 0 | 40.66 |
| Male_Medina_1 | m64468e_230405_211057.hifi_reads.fastq.gz | KAUST BCL | Sequel IIe | FASTQ | DNA | 1797853 | 38634465536 | 105 | 21489.2 | 50425 | 17311 | 20854 | 25053 | 0 | 22428 | 0 | 0 | 41.01 |
| Nabtat Ali | NabtatAli_rep1.ccs.fastq.gz | KAUST BCL | Sequel II | FASTQ | DNA | 1963722 | 32150514763 | 84 | 16372.2 | 49229 | 13366 | 15771 | 18827 | 0 | 16835 | 98.31 | 96.05 | 40.33 |
| Nabtat Ali | NabtatAli_rep2.ccs.fastq.gz | KAUST BCL | Sequel II | FASTQ | DNA | 2015273 | 33184869478 | 84 | 16466.7 | 49853 | 13435 | 15865 | 18951 | 0 | 16946 | 98.36 | 96.16 | 40.26 |
| Nabtat Saif | r64068_20220126_095227_4_D01.ccs.fastq.gz | KAUST BCL | Sequel II | FASTQ | DNA | 1114055 | 22678544018 | 84 | 20356.8 | 50039 | 16306 | 19557 | 23674 | 0 | 21106 | 98.1 | 95.49 | 40.31 |
| Nabtat Saif | r64068_20220126_095227_5_E01.ccs.fastq.gz | KAUST BCL | Sequel II | FASTQ | DNA | 1481192 | 29995850501 | 86 | 20251.2 | 50741 | 16215 | 19462 | 23583 | 0 | 21015 | 98.12 | 95.5 | 40.36 |
| Nabtat Sultan | m64468e_230205_130803.hifi_reads.fastq.gz | KAUST BCL | Sequel IIe | FASTQ | DNA | 1862512 | 34131352752 | 111 | 18325.4 | 50005 | 14814 | 17706 | 21248 | 0 | 19004 | 98.56 | 96.69 | 41.06 |
| Nabtat Sultan | m64468e_230212_132645.hifi_reads.fastq.gz | KAUST BCL | Sequel IIe | FASTQ | DNA | 1548427 | 27710398234 | 103 | 17895.8 | 49616 | 14585 | 17312 | 20625 | 0 | 18495 | 98.38 | 96.18 | 41.63 |
| Reziz | reziz_ccs.fastq.gz | KAUST BCL | Sequel II | FASTQ | DNA | 923581 | 17953944899 | 85 | 19439.5 | 50045 | 15900 | 18773 | 22395 | 0 | 20011 | 98.26 | 95.94 | 40.14 |
| Reziz | reziz_resequence_A01_ccs.fastq.gz | KAUST BCL | Sequel II | FASTQ | DNA | 788866 | 15202411733 | 86 | 19271.2 | 50172 | 15500 | 18498 | 22293 | 0 | 19904 | 98.18 | 95.65 | 40.49 |
| Rutana | Rotana_rep1.ccs.fastq.gz | KAUST BCL | Sequel II | FASTQ | DNA | 1867795 | 33865354767 | 109 | 18131.2 | 50277 | 14680 | 17589 | 21117 | 0 | 18888 | 98.56 | 96.62 | 40.42 |
| Rutana | Rotana_rep2.ccs.fastq.gz | KAUST BCL | Sequel II | FASTQ | DNA | 1680367 | 30260636813 | 105 | 18008.3 | 49694 | 14621 | 17458 | 20911 | 0 | 18717 | 98.33 | 96.06 | 40.57 |
| Safawi | Safawi_rep1.ccs.fastq.gz | KAUST BCL | Sequel II | FASTQ | DNA | 1440605 | 25605113553 | 124 | 17773.9 | 50021 | 14087 | 17125 | 20801 | 0 | 18588 | 98.31 | 96.03 | 40.68 |
| Safawi | Safawi_rep2.ccs.fastq.gz | KAUST BCL | Sequel II | FASTQ | DNA | 1661380 | 29851354178 | 126 | 17967.8 | 50236 | 14204 | 17317 | 21074 | 0 | 18826 | 98.52 | 96.58 | 40.68 |
| Sufri | m64468e_230206_221744.hifi_reads.fastq.gz | KAUST BCL | Sequel IIe | FASTQ | DNA | 1925231 | 34134293464 | 105 | 17730 | 50281 | 14615 | 17139 | 20331 | 0 | 18222 | 98.37 | 96.17 | 40.72 |
| Sufri | m64468e_230208_072500.hifi_reads.fastq.gz | KAUST BCL | Sequel IIe | FASTQ | DNA | 1787995 | 33773377643 | 154 | 18889 | 49727 | 15599 | 18343 | 21682 | 0 | 19461 | 98.6 | 96.8 | 41.31 |
| Sagai | Sakkai_rep1.ccs.fastq.gz | KAUST BCL | Sequel II | FASTQ | DNA | 1666793 | 28204024340 | 77 | 16921.1 | 48229 | 14050 | 16437 | 19396 | 0 | 17425 | 98.25 | 95.93 | 40.22 |
| Shahl | r64068_20220206_084143_2_B01.ccs.fastq.gz | KAUST BCL | Sequel II | FASTQ | DNA | 1376225 | 22683439403 | 86 | 16482.4 | 50039 | 13764 | 15898 | 18708 | 0 | 16776 | 98.42 | 96.33 | 40.52 |
| Shahl | r64068_20220206_084143_3_C01.ccs.fastq.gz | KAUST BCL | Sequel II | FASTQ | DNA | 1522318 | 25084569882 | 86 | 16477.9 | 50046 | 13761 | 15889 | 18699 | 0 | 16766 | 98.53 | 96.63 | 40.57 |
| Shishi | Shishi_rep1.ccs.fastq.gz | KAUST BCL | Sequel II | FASTQ | DNA | 2058023 | 36063130644 | 86 | 17523.2 | 49986 | 14291 | 16886 | 20183 | 0 | 18039 | 98.38 | 96.17 | 40.55 |
| Shishi | Shishi_rep2.ccs.fastq.gz | KAUST BCL | Sequel II | FASTQ | DNA | 2072726 | 35979183221 | 50 | 17358.4 | 49416 | 14158 | 16709 | 19970 | 0 | 17849 | 98.4 | 96.25 | 40.59 |
| Sukkari | Sukkary_rep1.ccs.fastq.gz | KAUST BCL | Sequel II | FASTQ | DNA | 1965313 | 32459809888 | 84 | 16516.4 | 50385 | 13598 | 15895 | 18892 | 0 | 16890 | 98.57 | 96.64 | 40.59 |
| Sukkari | Sukkary_rep2.ccs.fastq.gz | KAUST BCL | Sequel II | FASTQ | DNA | 2227392 | 37368207556 | 85 | 16776.7 | 50085 | 13818 | 16160 | 19207 | 0 | 17173 | 98.51 | 96.49 | 40.46 |
| Um Kabar | m64468e_230213_222801.hifi_reads.fastq.gz | KAUST BCL | Sequel IIe | FASTQ | DNA | 1432171 | 26710169961 | 118 | 18650.1 | 50133 | 14971 | 18015 | 21673 | 0 | 19375 | 0 | 0 | 42.05 |
| Um Kabar | m64468e_230215_073114.hifi_reads.fastq.gz | KAUST BCL | Sequel IIe | FASTQ | DNA | 1579378 | 28924439156 | 135 | 18313.8 | 50341 | 14734 | 17652 | 21205 | 0 | 18956 | 0 | 0 | 41.78 |

**Supplementary table 1.** a) Pacbio Sequencing Statistics generated with seqkit

| file | file names | Platform | | format | type | num_seqs | sum_len | min_len | avg_len | max_len | Q1 | Q2 | Q3 | sum_gap | N50 | Q20(%) | Q30(%) | GC(%) | coverage |
|---|---|---|---|---|---|---|---|---|---|---|---|---|---|---|---|---|---|---|---|
| Ajwa | DTG-OmniC-53_R1_001.fastq | Dovetail Genomics | Omni-C | FASTQ | DNA | 98523518 | 14778527700 | 150 | 150 | 150 | 150 | 150 | 150 | 0 | 150 | 97.22 | 93.02 | 44.79 | 42.8 |
| Ajwa | DTG-OmniC-53_R2_001.fastq | Dovetail Genomics | Omni-C | FASTQ | DNA | 98523518 | 14778527700 | 150 | 150 | 150 | 150 | 150 | 150 | 0 | 150 | 93.66 | 86.74 | 45.11 | |
| Anbara | AO5900046-pairsR1.fq.gz | Corteva Agriscience | HiC | FASTQ | DNA | 231131787 | 34669266218 | 24 | 150 | 150 | 150 | 150 | 150 | 0 | 150 | 97.22 | 92.88 | 41.12 | 100.5 |
| Anbara | AO5900046-pairsR2.fq.gz | Corteva Agriscience | HiC | FASTQ | DNA | 231131787 | 34668130570 | 24 | 150 | 150 | 150 | 150 | 150 | 0 | 150 | 94.79 | 88.25 | 41.77 | |
| Barni al-Ais | AO5900042-pairsR1.fq.gz | Corteva Agriscience | HiC | FASTQ | DNA | 121527508 | 18228862974 | 24 | 150 | 150 | 150 | 150 | 150 | 0 | 150 | 97.44 | 93.52 | 42.03 | 52.8 |
| Barni al-Ais | AO5900042-pairsR2.fq.gz | Corteva Agriscience | HiC | FASTQ | DNA | 121527508 | 18228277410 | 28 | 150 | 150 | 150 | 150 | 150 | 0 | 150 | 95.01 | 88.71 | 42.74 | |
| Barni al-Ula | AR2430006-pairsR1.fq.gz | Corteva Agriscience | HiC | FASTQ | DNA | 161247081 | 24186514148 | 80 | 150 | 150 | 150 | 150 | 150 | 0 | 150 | 96.05 | 89.76 | 42.85 | 70.1 |
| Barni al-Ula | AR2430006-pairsR2.fq.gz | Corteva Agriscience | HiC | FASTQ | DNA | 161247081 | 24185586423 | 24 | 150 | 150 | 150 | 150 | 150 | 0 | 150 | 95.97 | 89.94 | 43.27 | |
| Gurr | AR2430003-pairsR1.fq.gz | Corteva Agriscience | HiC | FASTQ | DNA | 104775995 | 15716033718 | 80 | 150 | 150 | 150 | 150 | 150 | 0 | 150 | 97.24 | 92.69 | 41.47 | 45.6 |
| Gurr | AR2430003-pairsR2.fq.gz | Corteva Agriscience | HiC | FASTQ | DNA | 104775995 | 15715439068 | 24 | 150 | 150 | 150 | 150 | 150 | 0 | 150 | 92.2 | 83.5 | 42.32 | |
| Hilali | AO5900045-pairsR1.fq.gz | Corteva Agriscience | HiC | FASTQ | DNA | 251098472 | 37664232940 | 24 | 150 | 150 | 150 | 150 | 150 | 0 | 150 | 97.45 | 93.28 | 41.6 | 109.2 |
| Hilali | AO5900045-pairsR2.fq.gz | Corteva Agriscience | HiC | FASTQ | DNA | 251098472 | 37663021013 | 24 | 150 | 150 | 150 | 150 | 150 | 0 | 150 | 94.78 | 88.12 | 42.04 | |
| Khudri | AO5900037-pairsR1.fq.gz | Corteva Agriscience | HiC | FASTQ | DNA | 106776223 | 16016329629 | 24 | 150 | 150 | 150 | 150 | 150 | 0 | 150 | 93.64 | 84.74 | 42.27 | 46.4 |
| Khudri | AO5900037-pairsR2.fq.gz | Corteva Agriscience | HiC | FASTQ | DNA | 106776223 | 16016096531 | 28 | 150 | 150 | 150 | 150 | 150 | 0 | 150 | 97.06 | 92.56 | 42.2 | |
| Male_Medina_1 | AR2430007-pairsR1.fq.gz | Corteva Agriscience | HiC | FASTQ | DNA | 116122075 | 17417891543 | 80 | 150 | 150 | 150 | 150 | 150 | 0 | 150 | 96.89 | 92.03 | 43.39 | 50.5 |
| Male_Medina_1 | AR2430007-pairsR2.fq.gz | Corteva Agriscience | HiC | FASTQ | DNA | 116122075 | 17417230916 | 24 | 150 | 150 | 150 | 150 | 150 | 0 | 150 | 94.74 | 87.77 | 43.99 | |
| Nabtat Ali | AO5900038-pairsR1.fq.gz | Corteva Agriscience | HiC | FASTQ | DNA | 110513305 | 16576757502 | 24 | 150 | 150 | 150 | 150 | 150 | 0 | 150 | 97.56 | 93.71 | 41.27 | 48.0 |
| Nabtat Ali | AO5900038-pairsR2.fq.gz | Corteva Agriscience | HiC | FASTQ | DNA | 110513305 | 16576228491 | 28 | 150 | 150 | 150 | 150 | 150 | 0 | 150 | 96.28 | 91.07 | 41.64 | |
| Nabtat Saif | AO5900036-pairsR1.fq.gz | Corteva Agriscience | HiC | FASTQ | DNA | 130718886 | 19607546020 | 24 | 150 | 150 | 150 | 150 | 150 | 0 | 150 | 97.34 | 93.27 | 41.87 | 56.8 |
| Nabtat Saif | AO5900036-pairsR2.fq.gz | Corteva Agriscience | HiC | FASTQ | DNA | 130718886 | 19606911452 | 24 | 150 | 150 | 150 | 150 | 150 | 0 | 150 | 96.44 | 91.67 | 42.33 | |
| Nabtat Sultan | AR2430004-pairsR1.fq.gz | Corteva Agriscience | HiC | FASTQ | DNA | 126409647 | 18960997343 | 80 | 150 | 150 | 150 | 150 | 150 | 0 | 150 | 97.29 | 93.06 | 43.87 | 55.0 |
| Nabtat Sultan | AR2430004-pairsR2.fq.gz | Corteva Agriscience | HiC | FASTQ | DNA | 126409647 | 18960290490 | 24 | 150 | 150 | 150 | 150 | 150 | 0 | 150 | 94.12 | 86.86 | 44.41 | |
| Reziz | AO4110006-pairsR1.fq.gz | Corteva Agriscience | HiC | FASTQ | DNA | 102485751 | 15371686699 | 24 | 150 | 150 | 150 | 150 | 150 | 0 | 150 | 85.33 | 76.37 | 39.35 | 44.6 |
| Reziz | AO4110006-pairsR2.fq.gz | Corteva Agriscience | HiC | FASTQ | DNA | 102485751 | 15474955270 | 24 | 151 | 151 | 151 | 151 | 151 | 0 | 151 | 85.68 | 76.22 | 39.38 | |
| Rutana | AO5900041-pairsR1.fq.gz | Corteva Agriscience | HiC | FASTQ | DNA | 243213241 | 36481464748 | 24 | 150 | 150 | 150 | 150 | 150 | 0 | 150 | 97.46 | 93.45 | 40.93 | 105.7 |
| Rutana | AO5900041-pairsR2.fq.gz | Corteva Agriscience | HiC | FASTQ | DNA | 243213241 | 36480319837 | 24 | 150 | 150 | 150 | 150 | 150 | 0 | 150 | 94.99 | 88.72 | 41.46 | |
| Safawi | AO5900039-pairsR1.fq.gz | Corteva Agriscience | HiC | FASTQ | DNA | 122265122 | 18339512356 | 24 | 150 | 150 | 150 | 150 | 150 | 0 | 150 | 97.25 | 92.98 | 41.56 | 53.2 |
| Safawi | AO5900039-pairsR2.fq.gz | Corteva Agriscience | HiC | FASTQ | DNA | 122265122 | 18338925759 | 24 | 150 | 150 | 150 | 150 | 150 | 0 | 150 | 95.79 | 90.22 | 41.97 | |
| Sufri | AR2430001-pairsR1.fq.gz | Corteva Agriscience | HiC | FASTQ | DNA | 131851909 | 19777323391 | 80 | 150 | 150 | 150 | 150 | 150 | 0 | 150 | 97.41 | 93.06 | 40.68 | 57.3 |
| Sufri | AR2430001-pairsR2.fq.gz | Corteva Agriscience | HiC | FASTQ | DNA | 131851909 | 19776583194 | 24 | 150 | 150 | 150 | 150 | 150 | 0 | 150 | 93.82 | 86.34 | 41.81 | |
| Sagai | AO5900040-pairsR1.fq.gz | Corteva Agriscience | HiC | FASTQ | DNA | 218170577 | 32725126833 | 24 | 150 | 150 | 150 | 150 | 150 | 0 | 150 | 97.36 | 93.18 | 41.37 | 94.9 |
| Sagai | AO5900040-pairsR2.fq.gz | Corteva Agriscience | HiC | FASTQ | DNA | 218170577 | 32724091200 | 24 | 150 | 150 | 150 | 150 | 150 | 0 | 150 | 95.79 | 90.12 | 41.81 | |
| Shahl | AO5900043-pairsR1.fq.gz | Corteva Agriscience | HiC | FASTQ | DNA | 138470101 | 20770228592 | 24 | 150 | 150 | 150 | 150 | 150 | 0 | 150 | 95.56 | 88.97 | 42.21 | 60.2 |
| Shahl | AO5900043-pairsR2.fq.gz | Corteva Agriscience | HiC | FASTQ | DNA | 138470101 | 20769581648 | 24 | 150 | 150 | 150 | 150 | 150 | 0 | 150 | 96.47 | 91.48 | 42.37 | |
| Shishi | AO5900035-pairsR1.fq.gz | Corteva Agriscience | HiC | FASTQ | DNA | 126219107 | 18932619969 | 24 | 150 | 150 | 150 | 150 | 150 | 0 | 150 | 93.88 | 85 | 42.06 | 54.9 |
| Shishi | AO5900035-pairsR2.fq.gz | Corteva Agriscience | HiC | FASTQ | DNA | 126219107 | 18932018175 | 24 | 150 | 150 | 150 | 150 | 150 | 0 | 150 | 95.37 | 88.87 | 41.82 | |
| Sukkari | AO5900035-pairsR1.fq.gz | Corteva Agriscience | HiC | FASTQ | DNA | 126219107 | 18932619969 | 24 | 150 | 150 | 150 | 150 | 150 | 0 | 150 | 93.88 | 85 | 42.06 | 54.9 |
| Sukkari | AO5900035-pairsR2.fq.gz | Corteva Agriscience | HiC | FASTQ | DNA | 126219107 | 18932018175 | 24 | 150 | 150 | 150 | 150 | 150 | 0 | 150 | 95.37 | 88.87 | 41.82 | |
| Um Kabar | AR2430005-pairsR1.fq.gz | Corteva Agriscience | HiC | FASTQ | DNA | 150699297 | 22604378788 | 80 | 150 | 150 | 150 | 150 | 150 | 0 | 150 | 96.7 | 91.34 | 42.16 | 65.5 |
| Um Kabar | AR2430005-pairsR2.fq.gz | Corteva Agriscience | HiC | FASTQ | DNA | 150699297 | 22603526318 | 24 | 150 | 150 | 150 | 150 | 150 | 0 | 150 | 94.98 | 88.09 | 42.68 | |

**Supplementary table 1.** b) HiC and Omni-C Sequencing Statistics generated with seqkit

| Assembly | Ajwa hap1 | Ajwa hap2 | Anbara | Barni al-Ais | Barni al-Ula | Gurr | Hilali | Khudri | ale Medina | Nabtat Ali | Nabtat Saif | abtat Sulta | Reziz | Rutana | Safawi | Sufri | Sagai | Shahl | Shishi | Sukkari | Um Kabar |
|---|---|---|---|---|---|---|---|---|---|---|---|---|---|---|---|---|---|---|---|---|---|
| **# contigs (>= 0 bp)** | 965 | 468 | 53 | 102 | 67 | 117 | 43 | 51 | 194 | 54 | 46 | 50 | 766 | 49 | 42 | 59 | 136 | 43 | 72 | 60 | 124 |
| **# contigs (>= 20 Kb)** | 951 | 464 | 53 | 102 | 66 | 117 | 43 | 51 | 194 | 53 | 46 | 50 | 764 | 49 | 42 | 59 | 136 | 43 | 72 | 60 | 121 |
| **# contigs (>= 50 Kb)** | 328 | 251 | 48 | 56 | 51 | 66 | 40 | 46 | 108 | 47 | 42 | 39 | 471 | 46 | 42 | 38 | 112 | 36 | 65 | 49 | 61 |
| **# contigs (>= 70 Kb)** | 161 | 153 | 47 | 46 | 44 | 57 | 37 | 45 | 58 | 43 | 39 | 31 | 247 | 45 | 39 | 34 | 96 | 34 | 62 | 47 | 55 |
| **# contigs (>= 100 Kb)** | 101 | 86 | 41 | 40 | 42 | 54 | 36 | 43 | 45 | 41 | 39 | 29 | 134 | 44 | 37 | 31 | 86 | 34 | 59 | 46 | 48 |
| **# contigs (>= 200 Kb)** | 70 | 53 | 38 | 32 | 36 | 46 | 35 | 40 | 39 | 34 | 35 | 28 | 59 | 36 | 35 | 30 | 66 | 33 | 51 | 39 | 43 |
| **# contigs (>= 500 Kb)** | 51 | 38 | 28 | 28 | 31 | 35 | 26 | 35 | 26 | 26 | 32 | 26 | 36 | 30 | 30 | 25 | 40 | 27 | 38 | 28 | 36 |
| **# contigs (>= 1 Mb)** | 38 | 27 | 27 | 27 | 24 | 30 | 22 | 30 | 24 | 20 | 26 | 23 | 26 | 25 | 25 | 24 | 27 | 23 | 27 | 24 | 32 |
| **# contigs (>= 2 Mb)** | 30 | 23 | 23 | 23 | 21 | 20 | 20 | 25 | 22 | 19 | 23 | 23 | 24 | 18 | 19 | 23 | 25 | 21 | 19 | 22 | 26 |
| **# contigs (>= 5 Mb)** | 24 | 20 | 22 | 20 | 18 | 19 | 19 | 21 | 20 | 19 | 20 | 22 | 23 | 18 | 19 | 22 | 23 | 19 | 18 | 21 | 22 |
| **# contigs (>= 10 Mb)** | 20 | 20 | 20 | 19 | 18 | 18 | 19 | 20 | 19 | 18 | 18 | 21 | 21 | 18 | 19 | 21 | 22 | 19 | 18 | 18 | 19 |
| **# contigs (>= 20 Mb)** | 17 | 18 | 19 | 18 | 17 | 18 | 19 | 18 | 18 | 18 | 18 | 18 | 18 | 18 | 18 | 19 | 16 | 18 | 18 | 18 | 18 |
| **Total length (>= 0 bp)** | 763098752 | 707197440 | 703128988 | 699925397 | 709323484 | 722622754 | 693603451 | 711706462 | 708825316 | 696248243 | 697414530 | 720374870 | 736178109 | 703569314 | 697545936 | 716186663 | 696749285 | 696289813 | 704819807 | 709779327 | 712822686 |
| **Total length (>= 20 Kb)** | 762844769 | 707128446 | 703128988 | 699925397 | 709307225 | 722622754 | 693603451 | 711706462 | 708825316 | 696230538 | 697414530 | 720374870 | 736139290 | 703569314 | 697545936 | 716186663 | 696749285 | 696289813 | 704819807 | 709779327 | 712767815 |
| **Total length (>= 50 Kb)** | 739811706 | 698703343 | 702974314 | 698407721 | 708799738 | 720920843 | 693499579 | 711541646 | 705265166 | 696044014 | 697276957 | 720034740 | 724416687 | 703465254 | 697545936 | 715393546 | 695915434 | 696066824 | 704537795 | 709372599 | 710758634 |
| **Total length (>= 70 Kb)** | **730388750** | **693126082** | **702915481** | **697820335** | **708423069** | **720432910** | **693324700** | **711487069** | **702386010** | **695812150** | **697107183** | **719566995** | **711429678** | **703398691** | **697371931** | **715152230** | **694979026** | **695951118** | **704371242** | **709249544** | **710418797** |
| **Total length (>= 100 Kb)** | 725383819 | 687701284 | 702415529 | 697359607 | 708253446 | 720195699 | 693247396 | 711296597 | 701331073 | 695637037 | 697107183 | 719397111 | 702216727 | 703315253 | 697207791 | 714901794 | 694141503 | 695951118 | 704105541 | 709165615 | 709844310 |
| **Total length (>= 200 Kb)** | 721139202 | 683243027 | 702065501 | 696355637 | 707440540 | 719039575 | 693099902 | 710839219 | 700489466 | 694602745 | 696507481 | 719225945 | 692402968 | 702170906 | 696906602 | 714709549 | 691188974 | 695823721 | 702869309 | 708355580 | 709134283 |
| **Total length (>= 500 Kb)** | 714894369 | 678572357 | 698804016 | 694657225 | 705554270 | 715235187 | 690410928 | 709281718 | 696375417 | 692492772 | 695760628 | 718512462 | 684985283 | 699882170 | 695334670 | 712974712 | 681506910 | 693727822 | 698253519 | 704441372 | 706902712 |
| **Total length (>= 1 Mb)** | 705614085 | 670925830 | 698076418 | 694102055 | 700474914 | 711640542 | 687663848 | 704971695 | 694860684 | 688223864 | 691508371 | 716433982 | 678254982 | 696968079 | 691533516 | 712224230 | 672741345 | 690820871 | 690122746 | 701498210 | 704185563 |
| **Total length (>= 2 Mb)** | 693774414 | 664061401 | 692117310 | 689017415 | 695490269 | 698520641 | 684459155 | 697860419 | 691441883 | 687203932 | 686841841 | 716433982 | 675684362 | 687032944 | 683418635 | 710919305 | 670419742 | 688357964 | 678912488 | 698690886 | 697049805 |
| **Total length (>= 5 Mb)** | 674545547 | 652946523 | 690101658 | 678611571 | 683837930 | 695693267 | 681192831 | 684469715 | 686762635 | 687203932 | 676975115 | 713617549 | 671796689 | 687032944 | 683418635 | 707630311 | 663195811 | 682434768 | 675007942 | 696041457 | 684577013 |
| **Total length (>= 10 Mb)** | 647856601 | 652946523 | 677525225 | 670773388 | 683837930 | 689660136 | 681192831 | 678710796 | 680785091 | 681474933 | 661136038 | 704595843 | 656186519 | 687032944 | 683418635 | 701697498 | 656001893 | 682434768 | 675007942 | 675784163 | 659507597 |
| **Total length (>= 20 Mb)** | 605828015 | 620425057 | 661201054 | 654126675 | 663869152 | 689660136 | 681192831 | 651642218 | 668455016 | 681474933 | 661136038 | 667503394 | 617604441 | 687032944 | 667125159 | 670294593 | 561649794 | 662563718 | 675007942 | 675784163 | 643366801 |
| **# contigs** | 965 | 468 | 53 | 102 | 67 | 117 | 43 | 51 | 194 | 54 | 46 | 50 | 766 | 49 | 42 | 59 | 136 | 43 | 72 | 60 | 124 |
| **Largest contig** | 71178219 | 59277250 | 60266042 | 77945624 | 77846521 | 80446534 | 79828620 | 78924758 | 77398804 | 78531549 | 70495270 | 78989254 | 60117163 | 78831913 | 79090776 | 60088354 | 59972322 | 60478471 | 78640515 | 76513229 | 76540041 |
| **Total length** | 763098752 | 707197440 | 703128988 | 699925397 | 709323484 | 722622754 | 693603451 | 711706462 | 708825316 | 696248243 | 697414530 | 720374870 | 736178109 | 703569314 | 697545936 | 716186663 | 696749285 | 696289813 | 704819807 | 709779327 | 712822686 |
| **GC (%)** | 42.39 | 41.22 | 41.53 | 41.21 | 41.44 | 41.84 | 41.02 | 41.68 | 41.05 | 40.99 | 41.3 | 41.94 | 41.04 | 41.22 | 41.13 | 41.79 | 41.28 | 41.14 | 41.6 | 41.28 | 41.66 |
| **N50** | **30899783** | **34143252** | **33924550** | **36846766** | **36633429** | **36812884** | **33945758** | **33923993** | **38793082** | **37201565** | **35434950** | **33915027** | **30982168** | **37448007** | **36533734** | **34564115** | **33872985** | **34797239** | **35983421** | **36338337** | **33461270** |
| **N90** | 2993222 | 16945824 | 20934805 | 20877220 | 24639080 | 21075070 | 24385073 | 20381426 | 20796342 | 24405782 | 20447130 | 20478717 | 9107851 | 24541917 | 20993504 | 21240896 | 16633630 | 21311992 | 20824763 | 20764785 | 20039179 |
| **auN** | 33227625.1 | 34447059.9 | 35991690 | 40012222.8 | 42305108.7 | 42404924.6 | 40677012.9 | 39215550.7 | 40972118.1 | 42933858.6 | 39960130 | 40056083.1 | 32685624.3 | 42957972.3 | 41175006.3 | 37223702.1 | 33902765.4 | 39467061.2 | 41566910.5 | 41612835.6 | 38251596.7 |
| **L50** | **9** | **8** | **9** | **7** | **7** | **7** | **8** | **8** | **7** | **7** | **7** | **8** | **9** | **7** | **7** | **8** | **8** | **8** | **7** | **7** | **8** |
| **L90** | 28 | 19 | 18 | 17 | 16 | 17 | 17 | 18 | 17 | 16 | 17 | 18 | 22 | 16 | 17 | 18 | 20 | 17 | 17 | 17 | 18 |
| **# N's per 100 kbp** | 0 | 0 | 0 | 0 | 0 | 0 | 0 | 0 | 0 | 0 | 0 | 0 | 0 | 0 | 0 | 0 | 0 | 0 | 0 | 0 | 0 |

**Supplementary table 2**. a) Assembly stastics of Ajwa phased haplotypes and 19 primary genome assemblies of the other date palm accessions

| Accession | Haplotype | Contigs | | | | | | | | | | Total length) | | | | | | | | | | Largest contig | Total length | GC (%) | N50 | N90 | auN | L50 | L90 |
|---|---|---|---|---|---|---|---|---|---|---|---|---|---|---|---|---|---|---|---|---|---|---|---|---|---|---|---|---|---|
| | | all | >= 20Kb | >= 50Kb | >= 70Kb | >= 100Kb | >= 200Kb | >= 500Kb | >= 1Mb | >= 2Mb | >= 5Mb | all | >= 20Kb | >= 50Kb | >= 70Kb | >= 100Kb | >= 200Kb | >= 500Kb | >= 1Mb | >= 2Mb | >= 5Mb | | | | | | | | |
| Ajwa | hap1 | 4068 | 4052 | 1367 | 439 | 172 | 71 | 51 | 38 | 30 | 24 | 915076931 | 914785769 | 809506221 | 756178698 | 734362091 | 721387199 | 714894369 | 705614085 | 693774414 | 674545547 | 71178219 | 915076931 | 42.11 | 29217884 | 47147 | 27718363 | 12 | 1657 |
| | hap2 | 1729 | 1725 | 991 | 417 | 149 | 53 | 38 | 27 | 23 | 20 | 780718640 | 780649646 | 750727471 | 717153581 | 695443256 | 683243027 | 678572357 | 670925830 | 664061401 | 652946523 | 59277250 | 780718640 | 41.16 | 30930021 | 86244 | 31209383.1 | 10 | 227 |
| Anbara | hap1 | 132 | 132 | 117 | 106 | 89 | 66 | 44 | 35 | 28 | 26 | 695477598 | 695477598 | 694882627 | 694227853 | 692854209 | 689623282 | 682055447 | 676291280 | 667083807 | 660981059 | 50281815 | 695477598 | 41.4 | 31343030 | 12176109 | 29273997 | 10 | 22 |
| | hap2 | 110 | 110 | 102 | 91 | 82 | 60 | 43 | 33 | 29 | 25 | 700439384 | 700439384 | 700096957 | 699477520 | 698736576 | 695471579 | 690020072 | 682949051 | 677306631 | 664582113 | 59624845 | 700439384 | 41.41 | 30904303 | 13716256 | 30487685.3 | 10 | 22 |
| Barni al-Ais | hap1 | 244 | 244 | 145 | 123 | 83 | 56 | 42 | 35 | 32 | 27 | 707458544 | 707458544 | 704132746 | 702837652 | 699509304 | 695747832 | 690874424 | 686289718 | 682530802 | 670586628 | 43089091 | 707458544 | 41.46 | 30565240 | 11885522 | 28038595.2 | 10 | 23 |
| | hap2 | 1389 | 1387 | 519 | 199 | 109 | 51 | 40 | 33 | 29 | 24 | 755829125 | 755790537 | 722427900 | 703935229 | 696730852 | 689185639 | 686039691 | 680895742 | 674668808 | 660851032 | 72921989 | 755829125 | 40.91 | 30309078 | 1251551 | 29450140.4 | 10 | 33 |
| Barni al-Ula | hap1 | 169 | 168 | 121 | 95 | 83 | 69 | 50 | 43 | 34 | 22 | 712635935 | 712621164 | 710900674 | 709379007 | 708406592 | 706426351 | 700343784 | 695401205 | 683292417 | 643536222 | 60790760 | 712635935 | 41.79 | 28996211 | 6516648 | 33807505.5 | 8 | 22 |
| | hap2 | 736 | 736 | 310 | 134 | 81 | 61 | 41 | 34 | 27 | 23 | 726010899 | 726010899 | 709126571 | 698957445 | 694691345 | 692210464 | 686184210 | 681116340 | 672233217 | 656050993 | 77318539 | 726010899 | 41.15 | 30573315 | 5236056 | 34536194.3 | 9 | 23 |
| Gurr | hap1 | 326 | 325 | 219 | 171 | 141 | 101 | 52 | 31 | 29 | 22 | 729534403 | 729515129 | 725553844 | 722701653 | 720234594 | 714438734 | 699164143 | 684360955 | 680511050 | 658658907 | 80016902 | 729534403 | 42.1 | 33603745 | 8102153 | 35229932.2 | 8 | 22 |
| | hap2 | 1287 | 1285 | 486 | 238 | 139 | 67 | 36 | 28 | 24 | 23 | 749724309 | 749686317 | 718473087 | 704255023 | 696215131 | 686975643 | 678065396 | 672631501 | 667365192 | 663468735 | 70273954 | 749724309 | 41.11 | 31030164 | 702408 | 34634662 | 8 | 31 |
| Hilali | hap1 | 128 | 128 | 109 | 88 | 70 | 55 | 46 | 38 | 30 | 25 | 690290804 | 690290804 | 689547850 | 688290666 | 686811836 | 684700309 | 681981600 | 675757018 | 663782727 | 647462635 | 55678820 | 690290804 | 41.11 | 31043846 | 8356779 | 30151378.9 | 9 | 22 |
| | hap2 | 104 | 104 | 92 | 81 | 77 | 60 | 44 | 37 | 29 | 24 | 691350674 | 691350674 | 690888264 | 690253837 | 689941349 | 687565107 | 681993993 | 677159418 | 666880401 | 652397359 | 78629200 | 691350674 | 41.15 | 30618064 | 11288744 | 34223241.6 | 9 | 20 |
| Khudri | hap1 | 227 | 227 | 145 | 109 | 86 | 68 | 49 | 37 | 31 | 23 | 708966785 | 708966785 | 705878375 | 703686918 | 701791074 | 699053167 | 693251833 | 684568067 | 676580647 | 650077449 | 76672404 | 708966785 | 41.51 | 31539597 | 7754450 | 32633033.4 | 9 | 21 |
| | hap2 | 136 | 136 | 124 | 111 | 101 | 86 | 47 | 29 | 28 | 23 | 706058527 | 706058527 | 705610075 | 704813692 | 703959719 | 701612037 | 689181859 | 676933096 | 675467208 | 659066534 | 77297364 | 706058527 | 41.89 | 29398583 | 10553414 | 35120430.6 | 8 | 21 |
| Male Medina 1 | hap1 | 237 | 237 | 172 | 117 | 84 | 57 | 48 | 40 | 34 | 27 | 703238404 | 703238404 | 700663777 | 697368503 | 694764246 | 691242477 | 688449630 | 682602580 | 673714458 | 649810347 | 75825240 | 703238404 | 41.34 | 30577566 | 8615582 | 31897494.8 | 9 | 25 |
| | hap2 | 1557 | 1557 | 939 | 418 | 194 | 69 | 38 | 34 | 28 | 25 | 796572590 | 796572590 | 770545047 | 740069093 | 721953123 | 705149822 | 696533645 | 693408793 | 683605433 | 673515533 | 59648988 | 796572590 | 41.07 | 29214838 | 119731 | 29472338.8 | 10 | 148 |
| Nabtat Ali | hap1 | 112 | 112 | 97 | 87 | 75 | 51 | 35 | 30 | 27 | 26 | 682660283 | 682660283 | 682105682 | 681526816 | 680507965 | 676956600 | 671910671 | 668259192 | 664460378 | 661692267 | 77009396 | 682660283 | 40.82 | 34155188 | 10778378 | 36191440.6 | 8 | 21 |
| | hap2 | 88 | 88 | 84 | 75 | 67 | 54 | 38 | 29 | 27 | 24 | 702382855 | 702382855 | 702220788 | 701679375 | 701032155 | 699205597 | 693814439 | 688008170 | 685685427 | 672680883 | 69217762 | 702382855 | 41.12 | 33571322 | 11138000 | 36345449.2 | 8 | 20 |
| Nabtat Saif | hap1 | 103 | 103 | 94 | 85 | 77 | 66 | 49 | 42 | 31 | 27 | 699439951 | 699439951 | 699100011 | 698552842 | 697895771 | 696195608 | 689691080 | 684889464 | 668214448 | 654699491 | 59308226 | 699439951 | 41.7 | 29985203 | 8070181 | 29843681.6 | 9 | 24 |
| | hap2 | 86 | 85 | 83 | 76 | 69 | 62 | 48 | 37 | 31 | 24 | 690156555 | 690136717 | 690055594 | 689611404 | 688973125 | 687961758 | 683687529 | 675650614 | 666714399 | 644098339 | 70124503 | 690156555 | 41.14 | 30798420 | 7177284 | 35248180.2 | 8 | 21 |
| Nabtat Sultan | hap1 | 1181 | 1179 | 493 | 218 | 119 | 69 | 49 | 39 | 34 | 27 | 759107408 | 759068952 | 731880412 | 715951352 | 707742406 | 700920924 | 694056714 | 686965869 | 680014038 | 659600253 | 77061567 | 759107408 | 41.28 | 28107417 | 1479084 | 29502104.6 | 10 | 36 |
| | hap2 | 161 | 161 | 113 | 93 | 73 | 57 | 43 | 34 | 29 | 26 | 718161315 | 718161315 | 716471116 | 715262929 | 713663145 | 711305358 | 707367727 | 701269160 | 694466568 | 682899879 | 59484578 | 718161315 | 41.91 | 31244978 | 10858331 | 32771845.3 | 9 | 21 |
| Reziz | hap1 | 979 | 976 | 597 | 365 | 209 | 136 | 99 | 81 | 67 | 49 | 740203417 | 740162987 | 725089195 | 711518327 | 698743740 | 689024581 | 678093519 | 665066481 | 645469888 | 586268844 | 35721589 | 740203417 | 40.78 | 11167579 | 895486 | 13622695.4 | 20 | 83 |
| | hap2 | 919 | 919 | 634 | 411 | 294 | 200 | 129 | 104 | 80 | 45 | 740450723 | 740450723 | 729204706 | 716413799 | 706611466 | 693747419 | 670103160 | 652294331 | 618568551 | 498660362 | 29558159 | 740450723 | 41.51 | 7737700 | 568876 | 10587071.9 | 25 | 123 |
| Rutana | hap1 | 98 | 98 | 90 | 80 | 73 | 49 | 32 | 28 | 24 | 21 | 688124011 | 688124011 | 687786208 | 687204242 | 686610543 | 683084786 | 677034816 | 674243965 | 667959772 | 654204128 | 54313506 | 688124011 | 41.01 | 33693003 | 20025634 | 34330828.5 | 8 | 19 |
| | hap2 | 101 | 101 | 93 | 85 | 80 | 59 | 43 | 33 | 27 | 23 | 697265524 | 697265524 | 696950562 | 696464445 | 696028232 | 693053598 | 687818558 | 681196435 | 671725746 | 656656739 | 76906267 | 697265524 | 41.37 | 31453009 | 10031318 | 35694862.7 | 8 | 20 |
| Safawi | hap1 | 91 | 91 | 86 | 83 | 78 | 67 | 52 | 43 | 32 | 28 | 698536044 | 698536044 | 698365880 | 698199480 | 697794878 | 696397659 | 691331144 | 685708393 | 670698889 | 656291844 | 59925122 | 698536044 | 41.53 | 31299995 | 9424216 | 29736679 | 9 | 24 |
| | hap2 | 95 | 95 | 88 | 84 | 73 | 54 | 39 | 29 | 27 | 24 | 692862607 | 692862607 | 692577699 | 692346894 | 691453877 | 688780448 | 684370903 | 676307250 | 673757104 | 665549333 | 77838405 | 692862607 | 41.16 | 33198872 | 11648730 | 36408517.4 | 8 | 19 |
| Sufri | hap1 | 150 | 150 | 108 | 78 | 66 | 55 | 44 | 37 | 31 | 28 | 712721270 | 712721270 | 711243674 | 709439149 | 708484441 | 707116735 | 703485844 | 698735305 | 691088056 | 683111816 | 59142343 | 712721270 | 41.75 | 30086201 | 10187603 | 28986234.4 | 10 | 23 |
| | hap2 | 1010 | 1010 | 410 | 178 | 88 | 48 | 38 | 33 | 28 | 20 | 743886522 | 743886522 | 720391980 | 707054864 | 699548972 | 694249525 | 691409817 | 687923501 | 680951480 | 653961435 | 78036640 | 743886522 | 41.16 | 30964179 | 3346195 | 34285482.3 | 9 | 24 |
| Sagai | hap1 | 319 | 319 | 246 | 209 | 176 | 122 | 74 | 60 | 50 | 40 | 702266027 | 702266027 | 699646200 | 697448559 | 694618456 | 687019598 | 672679952 | 662964540 | 648176191 | 616380124 | 34432499 | 702266027 | 41.57 | 18391943 | 3220294 | 17666139.4 | 14 | 45 |
| | hap2 | 216 | 216 | 199 | 171 | 157 | 118 | 68 | 56 | 49 | 41 | 698415396 | 698415396 | 697735067 | 696068926 | 694935444 | 689157892 | 673584526 | 665786358 | 655553880 | 629728474 | 33804412 | 698415396 | 41.36 | 19036978 | 5047761 | 17901185.9 | 15 | 41 |
| Shahl | hap1 | 123 | 123 | 105 | 87 | 79 | 60 | 45 | 39 | 33 | 31 | 685317483 | 685317483 | 684673662 | 683592654 | 682925895 | 680302770 | 675686632 | 671377369 | 662928929 | 656617109 | 55359055 | 685317483 | 41.08 | 25455583 | 9434020 | 27174209.3 | 10 | 26 |
| | hap2 | 84 | 84 | 77 | 71 | 63 | 57 | 47 | 38 | 32 | 26 | 696250550 | 696250550 | 695975432 | 695604788 | 694951491 | 693975435 | 691151332 | 685051289 | 675443766 | 658480938 | 56990679 | 696250550 | 41.26 | 29244976 | 9350505 | 31923431.4 | 9 | 22 |
| Shishi | hap1 | 160 | 160 | 134 | 117 | 100 | 77 | 48 | 32 | 27 | 24 | 695574537 | 695574537 | 694556005 | 693548609 | 692096849 | 688623253 | 679873646 | 669355931 | 662846346 | 649999768 | 60689813 | 695574537 | 41.38 | 29726859 | 9931187 | 34012633.7 | 8 | 22 |
| | hap2 | 204 | 204 | 185 | 163 | 145 | 88 | 43 | 30 | 24 | 21 | 711426892 | 711426892 | 710700516 | 709415301 | 707869624 | 699392984 | 685536808 | 676523962 | 668911258 | 658889137 | 70496341 | 711426892 | 41.86 | 32421388 | 12786191 | 37362815 | 8 | 19 |
| Sukkari | hap1 | 116 | 116 | 109 | 104 | 90 | 70 | 50 | 36 | 31 | 27 | 700206174 | 700206174 | 699941324 | 699645546 | 698530126 | 695542980 | 688317920 | 677988199 | 668949063 | 656539729 | 67437867 | 700206174 | 41.56 | 31696455 | 8125368 | 31887061.7 | 9 | 24 |
| | hap2 | 103 | 103 | 96 | 88 | 75 | 54 | 47 | 36 | 31 | 24 | 696818605 | 696818605 | 696530672 | 696053998 | 694952083 | 692029764 | 689972426 | 682384118 | 674648786 | 653614682 | 76353387 | 696818605 | 41.13 | 32905482 | 7012140 | 36846114.6 | 8 | 21 |
| Um Kabar | hap1 | 210 | 209 | 156 | 124 | 93 | 75 | 51 | 40 | 34 | 27 | 710179139 | 710159608 | 708159425 | 706246156 | 703715553 | 701305340 | 693320468 | 685670639 | 675626750 | 651516010 | 75641589 | 710179139 | 41.63 | 29970123 | 6094257 | 30120195.2 | 10 | 25 |
| | hap2 | 879 | 879 | 411 | 201 | 112 | 66 | 49 | 40 | 33 | 26 | 739479337 | 739479337 | 720846208 | 708622311 | 701282142 | 695341260 | 690099845 | 683802352 | 674418507 | 652265419 | 76083622 | 739479337 | 41.29 | 29320250 | 3396423 | 31510337.3 | 9 | 30 |

**Supplementary table 2**. b) Assembly stastics of haplotype assemblies of all 20 genome  date palm accessions a)

| genes | gene symbol | Chromosome | Haplotype | identity | length | mismatches | gaps open | q.start | q.end | s.start | s.end | e-value | bitscore |
|---|---|---|---|---|---|---|---|---|---|---|---|---|---|
| **CYP703** | LOC120107104 | Chr14 | Hap1 (male) | 100 | 1752 | 0 | 0 | 1 | 1752 | 3622873 | 3621122 | 0 | 3160 |
| **GPAT3** | LOC120108105 | Chr14 | Hap1 (male) | 99.76 | 12917 | 3 | 4 | 1 | 12894 | 9843968 | 9831057 | 0 | 23162 |
| **LOG3** | LOC103701078 | Chr16 | Hap1 (male) | 99.098 | 1885 | 5 | 6 | 3 | 1881 | 17953772 | 17955650 | 0 | 3307 |
| | | Chr16 | Hap2 (female) | 99.099 | 1886 | 4 | 7 | 3 | 1881 | 17575138 | 17577017 | 0 | 3305 |
| | | Chr14 | Hap1 (male) | 86.872 | 1912 | 127 | 27 | 15 | 1881 | 8456525 | 8458357 | 0 | 2307 |

b)

| gene | gene symbol | Chromosome | Accession | identity | length | mismatches | gaps open | q.start | q.end | s.start | s.end | e-value | bitscore |
|---|---|---|---|---|---|---|---|---|---|---|---|---|---|
| **LOG3** | LOC103701078 | Chr16 | Ajwa | 99.1 | 1885 | 5 | 6 | 3 | 1881 | 17754256 | 17756134 | 0 | 3307 |
| | | Chr16 | Anbara | 99.1 | 1885 | 5 | 6 | 3 | 1881 | 17826184 | 17828062 | 0 | 3307 |
| | | Chr16 | Barni al-Ais | 99.1 | 1885 | 6 | 6 | 3 | 1881 | 17770082 | 17771961 | 0 | 3306 |
| | | Chr16 | Barni al-Ula | 99.1 | 1886 | 4 | 7 | 3 | 1881 | 17735027 | 17736906 | 0 | 3305 |
| | | Chr16 | Gurr | 99.15 | 1886 | 4 | 6 | 3 | 1881 | 18004024 | 18005904 | 0 | 3314 |
| | | Chr16 | Hilali | 99.1 | 1887 | 4 | 6 | 3 | 1881 | 17703974 | 17705855 | 0 | 3312 |
| | | Chr16 | Khudri | 99.1 | 1886 | 4 | 7 | 3 | 1881 | 17974381 | 17976260 | 0 | 3305 |
| | | Chr16 (primary) | Male Medina 1 | 99.1 | 1885 | 5 | 6 | 3 | 1881 | 17953772 | 17955650 | 0 | 3307 |
| | | Chr14 (primary) | Male Medina 1 | 86.87 | 1912 | 127 | 27 | 15 | 1881 | 8456525 | 8458357 | 0 | 2307 |
| | | Chr16 | Nabtat Ali | 99.15 | 1887 | 5 | 6 | 1 | 1881 | 17788390 | 17790271 | 0 | 3314 |
| | | Chr16 | Nabtat Saif | 99.1 | 1886 | 4 | 7 | 3 | 1881 | 17985854 | 17987733 | 0 | 3305 |
| | | Chr16 | Nabtat Sultan | 99.1 | 1886 | 4 | 7 | 3 | 1881 | 17619385 | 17621264 | 0 | 3305 |
| | | Chr16 | Reziz | 99.1 | 1885 | 5 | 6 | 3 | 1881 | 17600835 | 17602713 | 0 | 3307 |
| | | Chr16 | Rutana | 99.1 | 1885 | 5 | 6 | 3 | 1881 | 17858337 | 17860215 | 0 | 3307 |
| | | Chr16 | Safawi | 99.1 | 1886 | 4 | 7 | 3 | 1881 | 17911086 | 17912965 | 0 | 3305 |
| | | Chr16 | Sufri | 99.1 | 1886 | 4 | 7 | 3 | 1881 | 17707224 | 17709103 | 0 | 3305 |
| | | Chr16 | Sakgi | 99.1 | 1887 | 4 | 6 | 3 | 1881 | 18213432 | 18215313 | 0 | 3312 |
| | | Chr16 | Shahl | 99.1 | 1885 | 5 | 6 | 3 | 1881 | 17972354 | 17974232 | 0 | 3307 |
| | | Chr16 | Shishi | 100 | 1881 | 0 | 0 | 1 | 1881 | 17907550 | 17909430 | 0 | 3393 |
| | | Chr16 | Sukkari | 99.1 | 1886 | 4 | 7 | 3 | 1881 | 18160254 | 18162133 | 0 | 3305 |
| | | Chr16 | Um Kabar | 99.1 | 1886 | 4 | 7 | 3 | 1881 | 17463207 | 17465086 | 0 | 3305 |

**Supplementary table 3.** Blasn results of 3 males specific genes indentified by Torres et al., (2019) on the male haplotypes (a) and all 20 date palm genomes (b).

**Supplementary Note**: **Comparison of the Ajwa genome assembly with previously released Khalas PDK50 genome**

As discussed in the main text, there are two date palm genome assemblies in the public domain that were generated with state-of-the-art technologies when released; however, these assemblies are not of the quality generated in our study. The female Khalas PDK50[1] genome was represented by 18 Linkage Groups (LGs), but contained an exceptionally high number of gaps (10,972). In contrast, the male Barhee BC4 genome[2], although having a much lower number of gaps (317), could not be fully anchored to the Khalas LGs, leaving ~50% of its contigs unplaced (i.e. 385 Mb out of 772 Mb).

As described in the main text, we generated a high-quality genome assembly for the Ajwa variety, successfully reconstructing all chromosomes with complete telomeric sequences and only a single gap in the Nucleolar Organizing Region (NOR). For a precise genome comparison, Ajwa chromosomes were aligned onto the Khalas genome using the scaffolding tool RagTag Scaffold [3,4]and the results were visualized using GS-Viewer (https://github.com/mirkocelii/GS-viewer). The Ajwa chromosomes exhibited substantial differences in size and showed only partial homology with the 18 linkage groups (LGs) of Khalas. In fact, only 52.73% of the Khalas LGs lengths could be covered by Ajwa homologous chromosome alignment (Supplementary Note Figures 1a-b). Two Ajwa chromosomes (Chr09 and Chr18) were scaffolded onto Khalas LG11, displaying a mosaic pattern of homology (Supplementary Note Figures 1 b-c). Unexpectedly, no Ajwa contigs could be scaffolded onto Khalas LG16 (Supplementary Note Figure 1a). To identify Ajwa sequences homologous to LG16, we generated a dotplot of all Ajwa chromosomes and all Khalas LGs, including unplaced contigs, using D-Genies [5]. The dotplot showed that Ajwa Chr01 was homologous to Khalas LGs 04 and 16, which suggested they may belong to a single chromosome (Supplementary Note Figure 1c).

Here we provide multiple evidence to support the Ajwa structure of Chr01, Chr09, and Chr18. First, as shown in the main text, the three Ajwa chromosomes are all gap-free and display telomeric sequences at both ends. Ajwa Chr01, ~~which is~~ homologous to the Khalas LG04 and LG16, matches a single optical map and displays enriched interactions across its entire sequence in the Hi-C contact maps. Ajwa Chr09 and Chr18, both homologous to LG11, correspond to two separate optical maps and did not display any enriched interactions in the Hi-C contact maps (Supplementary Note Figures 2-ab). Moreover, all other 19 high-quality genomes presented here exhibited the same structural features of Ajwa for chromosomes 01, 09, and 18 (Figure 2a,

Supplementary Note Figure 2d), and all are well supported by optical maps scaffolding (Supplementary File 2). Lastly, the Ajwa structures of Chr01, Chr09, and Chr18 were also consistent with their homologues in other palm species' genomes - e.g. coconut palm (*Cocos nucifera*[6]) and oil palms (*Elaeis oleifera*[7] and *Elaeis guineensis*[8]) (Figure 4).

To provide cytogenetic evidence of the proposed structure of the Ajwa genome, we designed a set of FISH probes homologous to Chr01, Chr09, and Chr18 (Supplementary Note Table 1).

Chromosome-specific oligonucleotide probes for Chr01, Chr09, and Chr18 were designed by Arbor Biosciences (Ann Arbor, MI) using the Ajwa genome assembly. Probe counts per target were as follows: 27392 probes for Chr01, 14812 probes for Chr09, and 9194 probes for Chr18. Each oligo probe set was labeled with either biotin-16-dUTP or digoxigenin-11-dUTP (Arbor Biosciences) and treated as for the 45S rDNA and Telomer probes. These probes were hybridized to Ajwa chromosome squashes (Supplementary Note Figure 2c) and confirmed that chromosomes 01, 09 and 18 were all separate molecules.

As previously mentioned, only 52.73% of the Khalas LG length is covered by each respective homologue Ajwa chromosome. We discovered that the remaining interspersed segments were instead aligning to the other Ajwa Chromosomes, creating a pattern of alternating homologous and non-homologous regions similar to the case of LG11 with Chr08 and Chr09 (Supplementary Note Figure 3a).

The anomalies between the assemblies presented in this study and the Khalas genome assembly can largely be attributed to the older technologies used over the past decade to sequence and assemble the Khalas genome (i.e. genetic mapping, first-generation long-read sequencing, and polishing with Illumina short reads), compared to the more advanced and recent approaches used in this study (i.e. PacBio HiFi reads, Hi-C, and optical mapping). However, we found that when Khalas LGs are split at gaps, and then re-scaffolded onto Ajwa with RagTag, a substantial improvement in homology and genome reconstruction could be observed (i.e. Ajwa coverage rises from 65.37% for the LGs, 82.71% for the split-contigs), except in highly repetitive putative centromeric and pericentromeric regions (Supplementary Note Figure 3b).

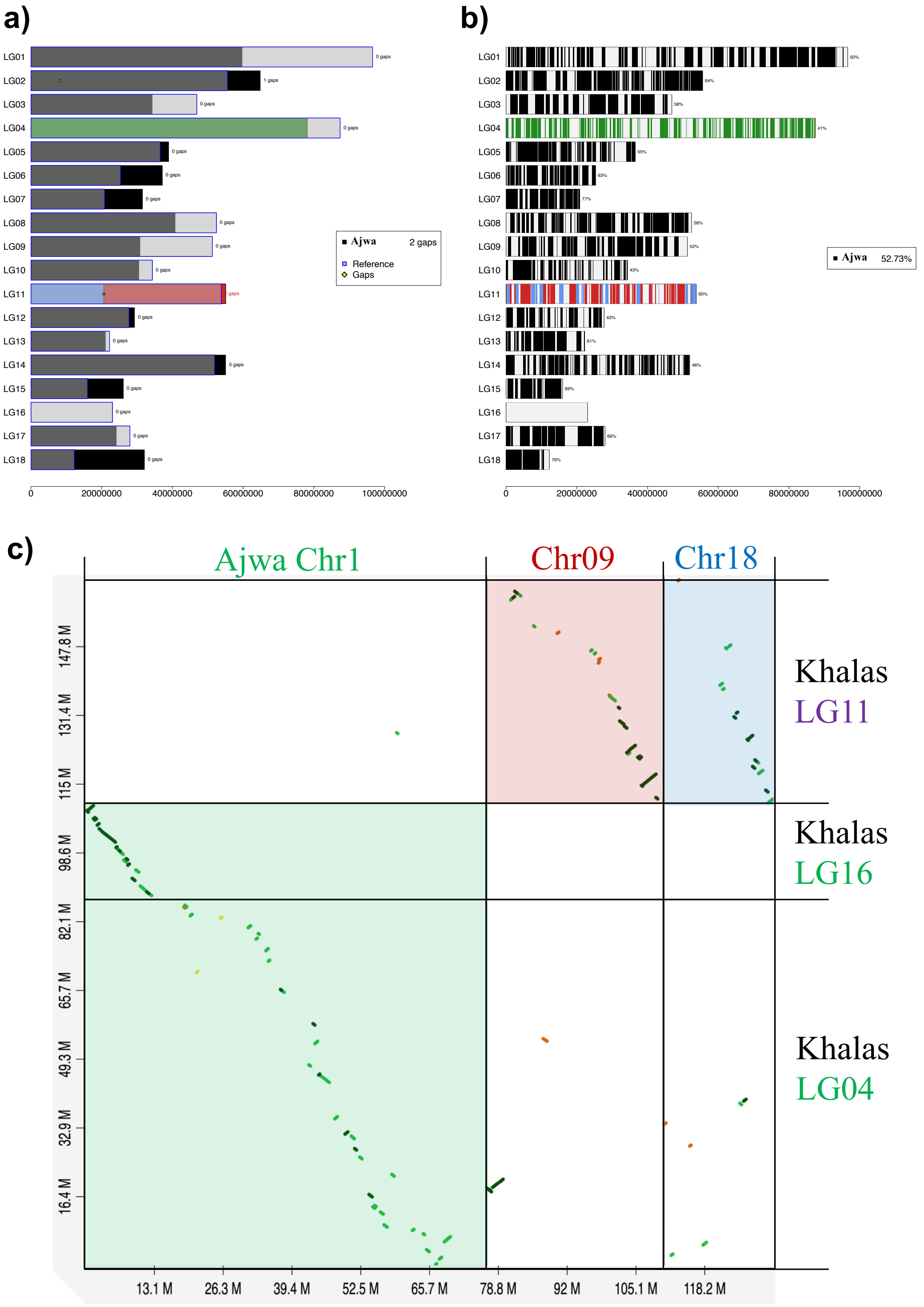


**Supplementary Note - Figure 1.**
a) Lengths of Ajwa chromosomes compared with the lengths of the Khalas linkage groups (LGs). b) Alignment of each Ajwa chromosome to its corresponding homologous Khalas LG visualized with GS-viewer scaffold. c) Dot plot generated with D-Genies[5] showing the alignment of Ajwa chromosomes Chr01, Chr09 and Chr18 with Khalas LG4, LG11 and LG16.

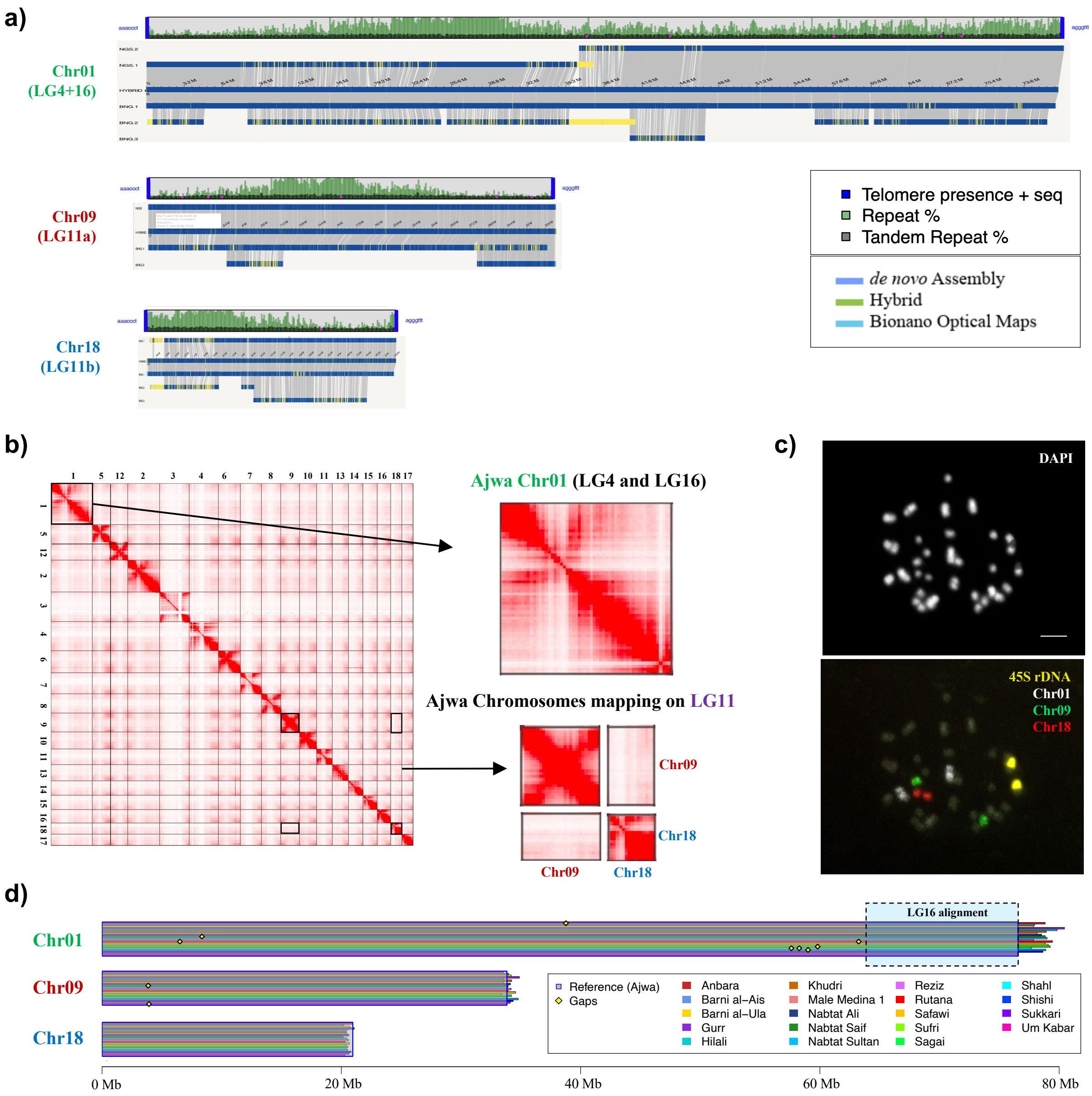


**Supplementary Note - Figure 2**

a) Repeat profiles of Chr01, Chr09 and Chr18 in 100-kb windows, overlaid with snapshots of Bionano hybrid-scaffold and optical maps.

b) Hi-C contact maps for Ajwa chromosomes, highlighting the cis interaction profile of Chr01 and the negligeable trans interactions between Chr09 and Chr18.

c) Fiber-FISH using probes designed specifically for Chr01, Chr09 and Chr18.

d) Length and gap positions of Chr01, Chr09 and Chr18 across 20 additional date palm genomes. The image was generated by aligning the 19 date palm genome assemblies to the Ajwa reference with RagTag and visualized with the GS-viewer scaffold.

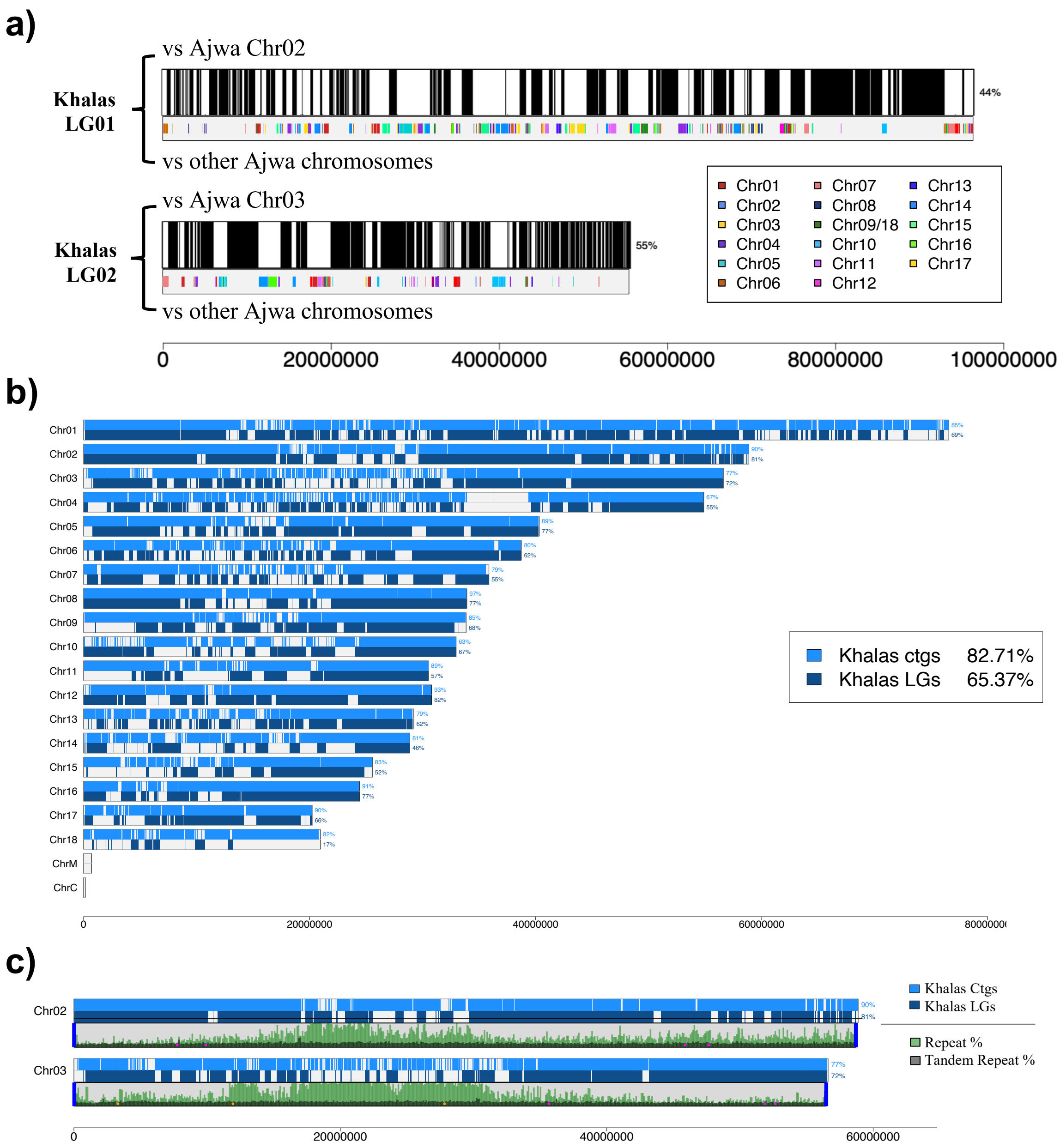


**Supplementary Note - Figure 3.**
a) Representation of Ajwa Chr02 and Chr03 aligned with RagTag to Khalas LG1 and LG2 and visualized with the GS-viewer scaffold module. The black track represents homologous alignments, whereas the multicoloured track represents non-homologous alignments, with one colour per chromosome.
b) Scaffolding of Khalas contigs and LGs to the Ajwa reference using RagTag and visualized with the GS-viewer scaffold module.
c) Overlay of the Khalas alignment profile on Ajwa Chr02 and Chr03 with the Ajwa repeat and tandem repeat profiles in 100-kb windows.
